# The reparative immunologic consequences of stem cell transplantation as a cellular therapy for refractory Crohn’s disease

**DOI:** 10.1101/2024.05.30.596699

**Authors:** Daniela Guisado, Sayali Talware, Xiaoli Wang, Andrew Davis, Elbek Fozilov, Aaron Etra, Jean-Frederic Colombel, Christoph Schaniel, Christopher Tastad, John E. Levine, James L. M. Ferrara, Ling-Shiang Chuang, Ksenija Sabic, Shishir Singh, Bridget K. Marcellino, Ronald Hoffman, Judy Cho, Louis J. Cohen

**Affiliations:** Division of Gastroenterology, Department of Medicine, Icahn School of Medicine at Mount Sinai, New York, NY, 10029, USA; Icahn Genomics Institute, Icahn School of Medicine at Mount Sinai, New York, NY, 10029, USA; Division of Pediatric Gastroenterology, Department of Pediatrics, Ichan School of Medicine at Mount Sinai, New York, NY, 10029, USA; Division of Hematology and Medical Oncology, Department of Medicine, Ichan School of Medicine at Mount Sinai, New York, NY, 10029, USA; Tisch Cancer Institute, Ichan School of Medicine at Mount Sinai, New York, NY, 10029, USA; Institute for Regenerative Medicine, Ichan School of Medicine at Mount Sinai, New York, NY, 10029, USA; Black Family Stem Cell Institute, Ichan School of Medicine at Mount Sinai, New York, NY, 10029, USA; Department of Pathology, Molecular and Cell Based Medicine, Icahn School of Medicine at Mount Sinai, New York, NY, 10029, USA; The Charles Bronfman Institute for Personalized Medicine, Icahn School of Medicine at Mount Sinai, New York, NY, 10029, USA

**Keywords:** Crohn’s Disease, Hematopoietic stem cells, Stem cell transplant, Macrophage, Autoimmune disease

## Abstract

**Background:** Treatment strategies for Crohn’s disease (CD) suppress diverse inflammatory pathways but many patients remain refractory to treatment. Autologous hematopoietic stem cell transplantation (SCT) has emerged as a therapy for medically refractory CD. SCT was developed to rescue cancer patients from myelosuppressive chemotherapy but its use for CD and other immune diseases necessitates reimagining SCT as a cellular therapy that restores appropriately responsive immune cell populations from hematopoietic progenitors in the stem cell autograft (i.e. immune “reset”). Here we present a paradigm to understand SCT as a cellular therapy for immune diseases and reveal how SCT re-establishes cellular immunity utilizing high-dimensional cellular phenotyping and functional studies of the stem cell grafts.

**Methods:** Immunophenotyping using CyTOF, single cell RNA sequencing (scRNA-seq) and T cell receptor (TCR) sequencing was performed on peripheral blood and intestinal tissue samples from refractory CD patients who underwent SCT. The stem cell graft from these patients was analyzed using flow cytometry and functionally interrogated using a murine model for engraftment.

**Results:** Our study revealed a remodeling of intestinal macrophages capable of supporting mucosal healing that was independently validated using multimodal studies of immune reconstitution events including CyTOF and scRNA-seq. Functional interrogation of hematopoietic stem cells (HSCs) using a xenograft model demonstrated that HSCs shape the timing of immune reconstitution, the selected reconstitution of specific cell lineages and potentially the clinical efficacy of SCT.

**Conclusions:** These studies indicate that SCT serves as a myeloid-directed cellular therapy re-establishing homeostatic intestinal macrophages that support intestinal healing and suggest refractory CD evolves from impairment of restorative functions in myeloid cells. Furthermore, we report heterogeneity among HSCs from CD patients which may drive SCT outcomes and suggests an unrecognized impact of CD pathophysiology on HSC in the marrow niche.

## INTRODUCTION

Crohn’s disease (CD) is an immune disease that involves dysregulation of host-microbe interactions causing inflammation of the gastrointestinal tract.^1^ The standard treatment strategy for CD targets inflammatory pathways. Despite an increasing diversity of immunotherapeutic strategies there remain a large number of patients who lose their initial therapeutic response increasing patient morbidity and mortality.^2^ The acquisition of a medically refractory phenotype appears independent of therapeutic mechanism as failure of any therapeutic modality leads to diminished efficacy with subsequent modalities. As such the development of medically refractory CD may be due to a redundancy in seemingly diverse immunotherapeutic strategies, or that causes of medically refractory CD cannot be corrected by suppression of inflammatory pathways alone. Autologous hematopoietic stem cell transplantation (SCT) is an emerging therapeutic option for patients with medically refractory CD as well as other immune diseases.^3^ The clinical efficacy of SCT in the majority of medically refractory CD patients presents an opportunity to understand novel therapeutic pathways by which SCT reverses the processes underlying refractory CD.^3–7^

SCT was developed for the treatment of cancer patients to facilitate the administration of high dose chemotherapy by utilizing hematopoietic stem cells (HSCs) to prevent chemotherapy induced myelosuppression. It is hypothesized that as a treatment for immune diseases SCT functions as a cellular therapy restoring appropriately responsive immune cell populations from hematopoietic progenitors in the stem cell autograft (i.e. an immune “reset”).^3^ Studies in patients with cancer have examined immune reconstitution following SCT but have not systematically explored such events as they might be relevant to SCT as a cellular therapy including how HSC subpopulations re-establish cellular immunity in tissue niches.^8–10^ Initial studies to understand mechanisms by which SCT alleviates the consequences of medically refractory CD profiled circulating immune cells using flow cytometry, but these studies did not shed light on how HSCs drive immune reconstitution events in the intestine.^8, 11^ Here we outline a paradigm to understand SCT as a cellular therapy for immune diseases and report our observations of how SCT determines immune reconstitution events in CD using high dimensional cellular phenotyping and functional studies of the stem cell grafts. Using single cell RNA sequencing (i.e. scRNA-seq) as well as mass cytometry (i.e. CyTOF) we have characterized immunophenotypic changes in the peripheral blood and the intestine of recipient patients that indicates SCT primarily acts as a myeloid-directed cellular therapy restoring intestinal macrophages capable of supporting mucosal healing. These studies highlight the central role of monocytes/macrophages in driving CD pathophysiology and propose the acquisition of medically refractory CD may not be secondary to unknown inflammatory pathways but rather the impairment of pathways that support mucosal healing. Functional studies of stem cell grafts in a xenograft model further revealed a previously unrecognized heterogeneity in the composition and function of mobilized HSCs which may determine these immune reconstitution events, SCT efficacy and suggest that CD pathophysiology extends to the marrow niche.

## METHODS

### MASCT-CD trial

Maintenance in Autologous Stem Cell Transplant for Crohn’s Disease (MASCT-CD) trial is a Phase 2 trial conducted at the Mount Sinai Hospital (New York, NY) with IRB approval through Mount Sinai (ClinicalTrials.gov NCT03219359, IRB study #16-01304). Adults with CD were eligible for SCT if they had active clinical and endoscopic disease and failed to respond to at least one member of each FDA approved drug class (anti-TNFα, anti-α4β7, anti-IL12/23), an immunomodulator (6MP, AZA, MTX) and if the patient was not a candidate for intestinal surgery. Patients underwent SCT per published protocols and were placed on vedolizumab at standard dosing immediately following hematological recovery.^4, 6^

### CyTOF

Isolated cells were processed and acquired on a CyTOF2 Mass Cytometer. CyTOF FCS files were concatenated and normalized using the bead-based normalization tool in the Helios software (Fluidigm), the barcoded samples were automatically deconvoluted and cross-sample doublets were filtered using a Matlab-based debarcoding tool^12^ and normalized and analyzed with Cytobank (Beckman Coulter). Supervised analysis was performed using cell population identification with known surface markers and previously established gating strategy.^13^ FlowSOM and CITRUS analyses were performed for unsupervised analyses and viSNE plots were created with concatenated samples.

### Xenograft model

NOD.Cg-*Prkdc^scid^ Il2rg^tm1Wjl^*/SzJ (NSG) mice were purchased from Jackson Laboratory (Bar Harbor, Maine). All experiments were approved by the Institutional Animal Care Committee of the Icahn School of Medicine at Mount Sinai (Protocol #202200000018) and followed strict IACUC guidelines. All mice were 8-9 week-old females and irradiated prior to transplant (220cGy). Thawed patient mobilized mononuclear cells were CD34+ selected using magnetic bead separation and 0.2-0.4×10^6^ CD34^+^ cells were transplanted via the tail vein into the irradiated mice. Mice were sacrificed at 4, 8, and 16 weeks after transplantation and cells were harvested from the peripheral blood, bone marrow, and spleen. We considered human engraftment to have occurred if human CD45^+^ cells were present at ≥0.1% of the nucleated cells in the marrow or spleen.

### Single Cell Data Processing

FASTQ files underwent processing through the Cellranger Count v6.1.1 pipeline for quality control, alignment, and matrix generation. The filtered matrices were employed in the Seurat wrapper coreSC for data cleaning and regularization and a Seurat object was created. Batch effects were addressed using the Harmony package. Differentially expressed genes were identified and used for annotation and cluster comparisons. We utilized dotplot functions, RNA velocity analysis using scVelo, and CellphoneDB for additional analyses.

### Statistical analyses

Data are reported as mean ± standard error of mean (SEM). Non-parametric tests including Mann-Whtiney and Wilcoxon tests were primarily used for analyses. Statistical analysis was performed using Prism 10 (GraphPad Software), R for scRNA-seq, and ImmunoSEQ Analyzer (Adaptive Biotechnologies) for TCR data. A P value of <0.05 was considered statistically significant.

See supplementary methods for more details.

## RESULTS

### SCT is highly effective in refractory CD patients

Patients with medically refractory CD were enrolled in the MASCT-CD trial. Nineteen patients were enrolled from 2018-2023 and 14 were followed for at least 6 months follow up post-SCT (Figure 1A). At 6 months post-transplant 10/14 patients had an endoscopic remission (Total SES-CD ≤ 4 and no sub-score >1) and 13/14 had endoscopic response (reduction of SES-CD by at least 50%) (Figure 1B-D).^14–16^ Paired blood and intestinal tissue samples were obtained at baseline and 6 months post-SCT and additional blood samples were taken longitudinally (Figure 1E).

**Figure 1.**
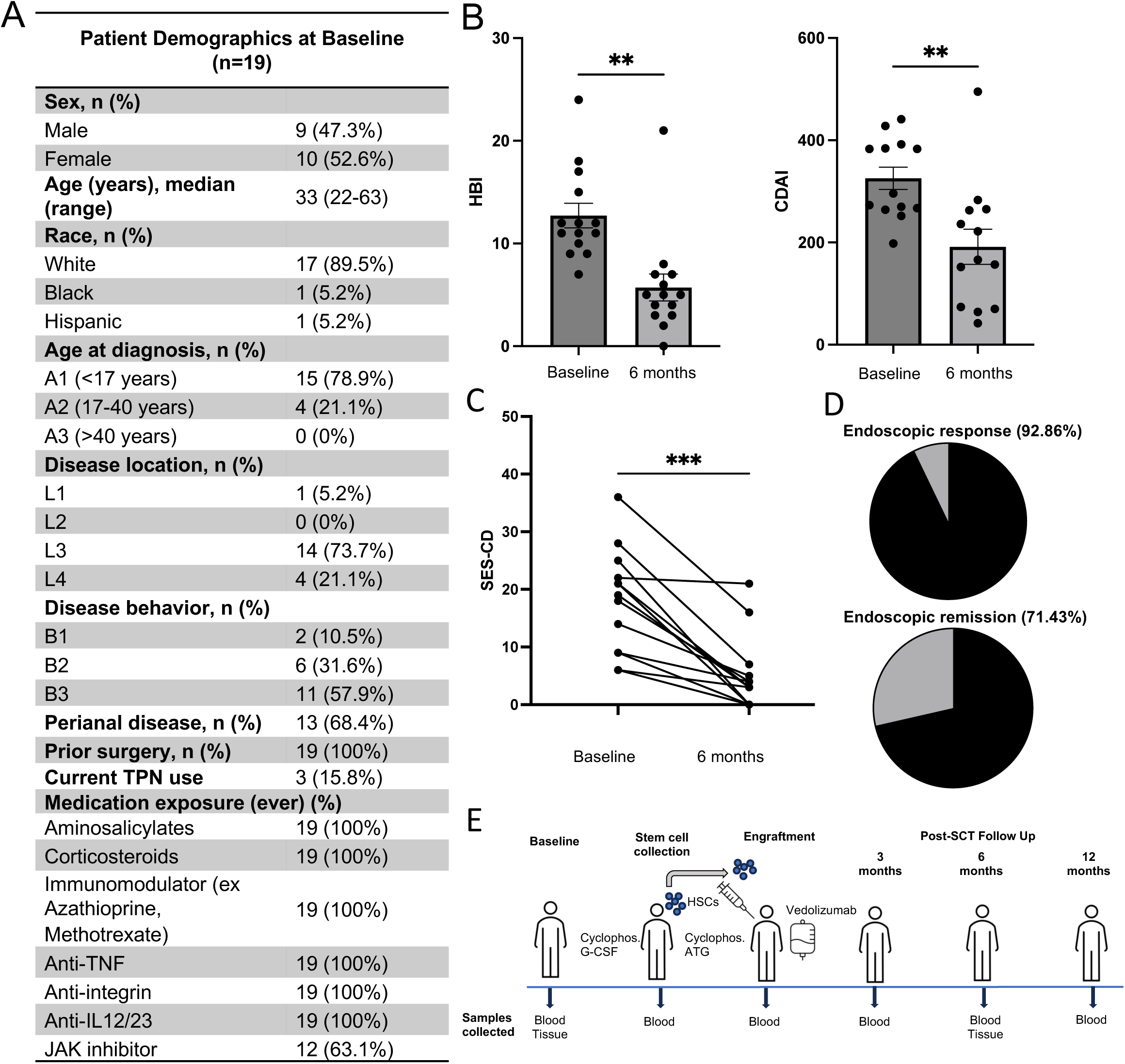
SCT is highly effective in refractory CD patients. **(A**) Table of patient demographics at baseline. **(B)** Bar graphs of clinical outcomes at 6 months post-SCT for HBI and CDAI scores (*n*=14, mean ± SEM). **(C)** Line graph of paired endoscopic outcomes with SES-CD at 6 months post-SCT (*n*=14). **(D)** Pie graphs of endoscopic response (SES-CD decrease ≥ 50%) and endoscopic remission (SES-CD ≤ 4 and no subscore >1) at 6 months post-SCT. All statistical analysis (B-C) done using paired Wilcoxon signed-rank test with * P<0.05, ** P<0.01 *** P<0.001 **** P<0.0001. **(E)** Schematic of MASCT-CD clinical trial and sample collection timeline.

### SCT has a distinct effect on intestinal immune populations

To characterize immune reconstitution events following SCT, blood and intestinal samples were analyzed longitudinally by mass cytometry (i.e. CyTOF). Supervised clustering of immune cell populations was performed using canonical markers and unsupervised clustering using FlowSOM.^13,17^ Iterative FlowSOM analyses performed independently on intestinal and blood samples organized immune cells into 15 metaclusters (i.e. parental population) inclusive of 100 cell clusters (i.e. sub-population). Metaclusters demonstrated concordance with supervised immune cell populations (Supplementary Table 1).

The effect of SCT on immune cell populations 6 months post-SCT suggested distinct changes to the circulating and the intestinal immune cell compartments (Figure 2A, Supplementary Figure 1). Analysis of FlowSOM organized metaclusters identified significant changes to circulating lymphoid populations and intestinal myeloid populations reinforcing the specific effect of the transplant on each immune compartment. Analysis of metacluster sub-populations (i.e. clusters) indicated far greater changes to intestinal immune cell populations with significant changes in 21/100 immune cell clusters compared to circulating immune cell clusters (3/100 clusters) (Figure 2B). A principal component analysis of FlowSOM organized immune cell clusters supported an effect of SCT on intestinal immune cell populations (Figure 2B). Supervised and unsupervised analyses demonstrated that the most significant changes post-SCT involved circulating CD27^-^ B cells (metacluster 3) and intestinal CD14^+^ cells (metacluster 1) (Figure 2C, Supplementary Figure 1). Prior studies of immune reconstitution events post-SCT in patients with CD similarly reported an effect on circulating naïve B cell populations.^8^ Our findings reveal a distinct and significant effect of SCT on intestinal immune cell populations highlighting the importance of defining immune reconstitution events in different niches including blood and intestine.

**Figure 2.**
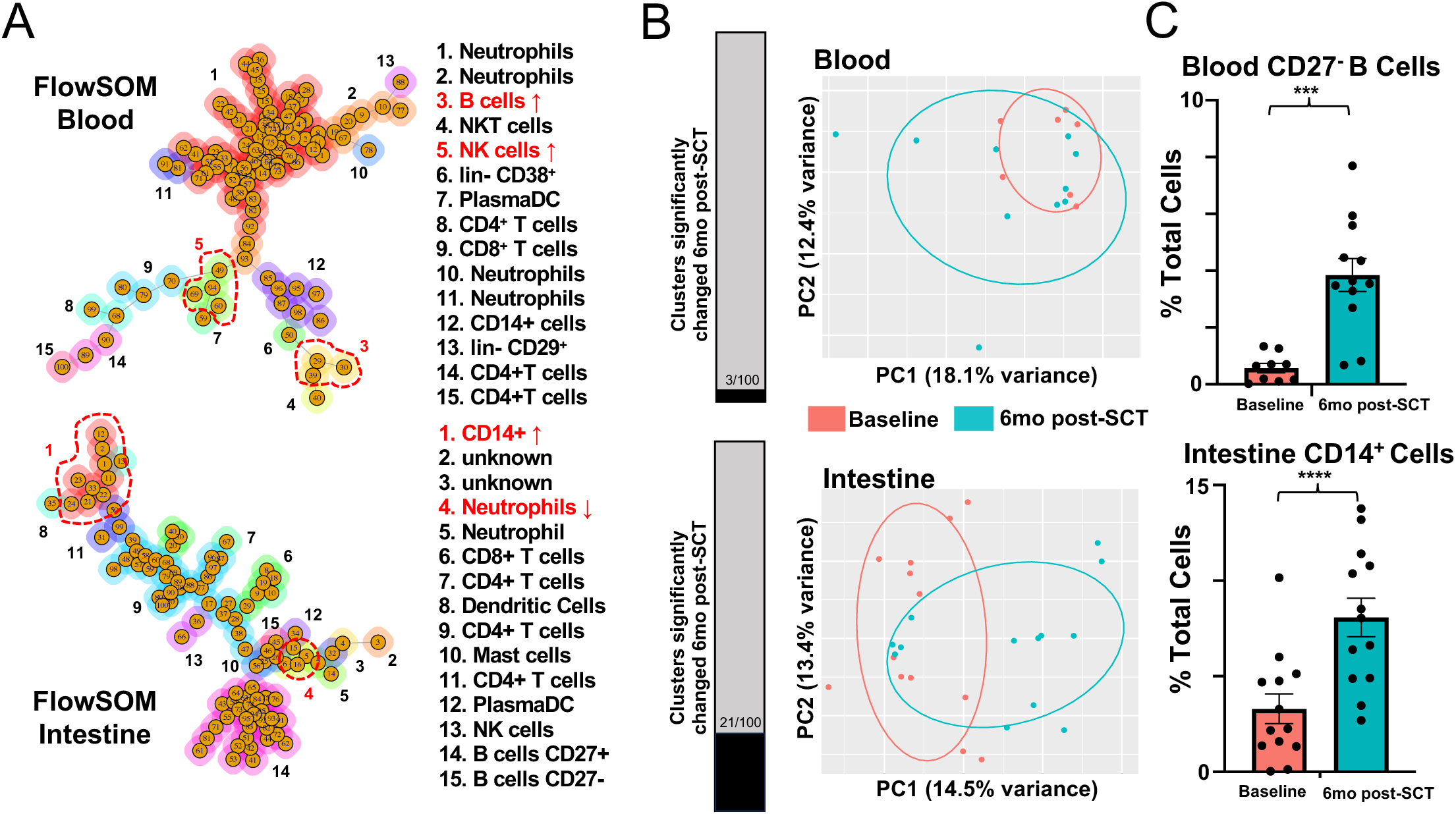
SCT has a distinct effect on intestinal immune populations. **(A**) Unsupervised CyTOF analysis with FlowSOM for blood (top) (baseline *n*=7, 6 months post-SCT *n*=9) and intestinal (bottom) (baseline *n* =13, 6 months post-SCT *n*=13) samples with each minimal spanning tree diagram displaying 15 metaclusters comprised of 100 clusters. Metaclusters annotated using canonical markers or prominent markers if lineage markers are negative (lin-CD3^-^ CD19^-^CD14^-^CD16^-^CD56^-^CD66b^-^). Metaclusters that were significantly changed from baseline to 6 months post-SCT are indicated by red font and red dashed circle (P<0.05, Mann-Whitney test). **(B)** Bar graph depicts the number of significantly changed clusters from baseline to 6 months post-SCT (P<0.05, Mann-Whitney test). Principal component analysis (PCA) of blood (top) and intestine (bottom) samples comparing baseline with 6 months post-SCT (FlowSOM clusters, PCA analysis 95% confidence interval). **(C)** Bar graph (mean ± SEM) of most significantly changed cell populations in supervised clustering analyses from blood (baseline *n*=9, 6 months post-SCT *n*=11) and intestine (baseline *n*=13, 6 months post-SCT *n*=13), Mann-Whitney test, *** P<0.001, **** P<0.0001.

### Reconstitution of intestinal myeloid populations

To further understand how immune sub-populations are collectively reconstituted post-SCT, CyTOF datasets were analyzed by CITRUS.^18^ CITRUS organizes single cells into hierarchical clusters that predict how a system responds to a perturbation (e.g. SCT). CITRUS analysis confirmed that a larger number of intestinal immune cell clusters were affected by SCT as compared to circulating immune cell clusters (Figure 3A and 3B). Among circulating immune cells there was a significant increase post-SCT in a single cluster network including 3 related transitional B cell clusters that are CD19^+^CD27^-^CD38^+^BTLA^+^ (Figure 3A, Supplementary Figure 7).^19, 20^ These transitional B cell clusters expressed HLADR, CD45RA, CD1c, CCR6 and integrins β7 and CD49d (α4) suggesting intestinal migrating transitional B cell populations including memory cells.^21^ Parental cell clusters to the B cell clusters did not express canonical lineage markers suggesting that they were progenitor cell populations. These progenitor cell clusters expressed integrins CD62L, CD29 and CD49d(α4) consistent with their potential to participate in intestinal trafficking.^22, 23^

**Figure 3.**
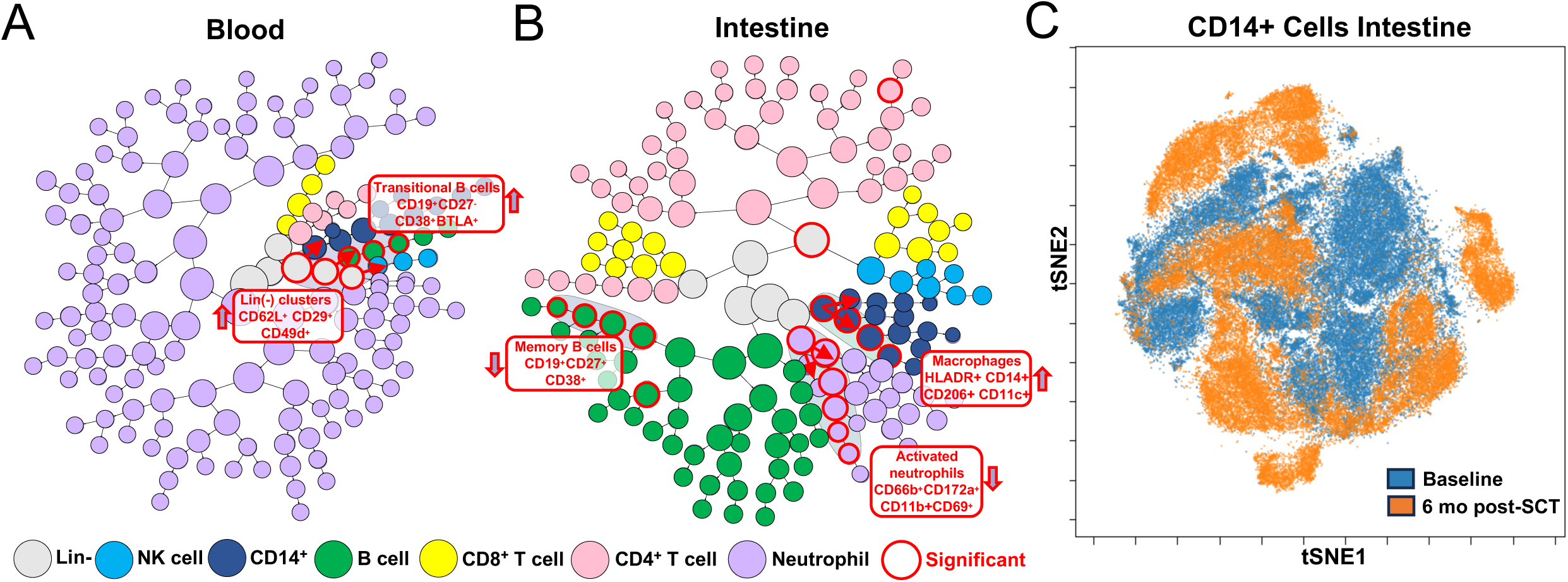
Reconstitution of intestinal myeloid populations. **(A)** CITRUS analysis of CyTOF datasets from blood (baseline *n*=9, 6 months post-SCT *n*=11) and **(B)** intestinal (baseline *n*=13, 6 months post-SCT *n*=13) samples. Hierarchical clusters are depicted as a CITRUS tree with each node representing a different cluster colored coded by immune cell populations based on canonical protein markers. Cluster networks (>1 cluster) that collectively predict the change in immune cell populations 6 months post-SCT compared to baseline are outlined in red (SAM, FDR <0.01). Hierarchical relationships between significant clusters are identified with red arrows indicating direction from parental to child cluster. **(C)** CD14+ population from CyTOF samples concatenated at each timepoint analyzed by viSNE using tSNE plots.

CITRUS analysis of the intestinal immune compartment identified significant changes post-SCT to three large cell networks (Figure 3B). These three cell networks included B cells, neutrophils and CD14^+^ cells (Supplementary Figure 8).^24^ In contrast to the circulating B cell network there are a small number of CD19^+^CD27^+^CD38^+^ intestinal memory B cell clusters which were significantly decreased post-SCT. A neutrophil cell network was also significantly reduced post-transplantation. These neutrophil clusters expressed traditional markers including CD66b, CD11c, CD33, CD172a, and CD11b.^25^ Neutrophil cell clusters that were significantly decreased post-SCT were distinct from neutrophil clusters unaffected by SCT based on their expression of CD69 suggesting a specific loss post-SCT of activated neutrophils.^26^ The CD14^+^ cell cluster network significantly increased post-SCT. This CD14^+^ cell cluster network expressed macrophage markers including HLADR, CD33, CD206, CD64, CD11b, CD11c and CD4.^27^ Within this CD14^+^ cell network were clusters that variably expressed CD206, CD11b and CD11c delineating changes to early differentiating monocytes/macrophages (CD11c^hi^) as well as mature macrophage populations (CD206^hi^CD11b^lo^).^27^ For each major myeloid population (e.g. neutrophil, CD14^+^ cells) there was a significant change in the parental cluster for the lineage suggesting that the SCT perturbs intestinal myeloid lineages (Figure 3B). To visualize global effects of SCT on major intestinal cell lineages we performed a viSNE analysis of manually gated T cell, B cell, neutrophil and CD14^+^ cell populations. This analysis confirmed dramatic effects of SCT on intestinal myeloid populations with almost complete phenotypic remodeling of CD14^+^ cells consistent with reconstitution of the myeloid lineage post-SCT (Figure 3C, Supplementary Figure 2).

### SCT restores homeostatic macrophages in refractory CD patients

Proteomic (i.e CyTOF) studies suggested a disproportionate effect of SCT on intestinal myeloid populations. To further explore functional changes among intestinal cell populations, samples at baseline and 6 months post-SCT were analyzed with scRNA-seq. Clustered scRNA-seq data were analyzed by CellphoneDB to define ligand-receptor pairs that comprised cellular interaction networks (Figure 4A, Supplementary Table 2).^28^ Among cell clusters, the stromal, myeloid, endothelial and glial cell clusters were predicted to interact extensively in the intestine whereas lymphoid clusters (B cell, T cell) had the smallest predicted interaction networks (Figure 4B). CellphoneDB identified the strongest interaction network involving myeloid cluster 17, stromal cluster 19 and glial cluster 21 (i.e. hub clusters) (Figure 4A). Hub clusters interacted broadly with other intestinal cell clusters confirming the importance of myeloid, stromal and glial cells as regulators of intestinal homeostasis.^29–32^

**Figure 4.**
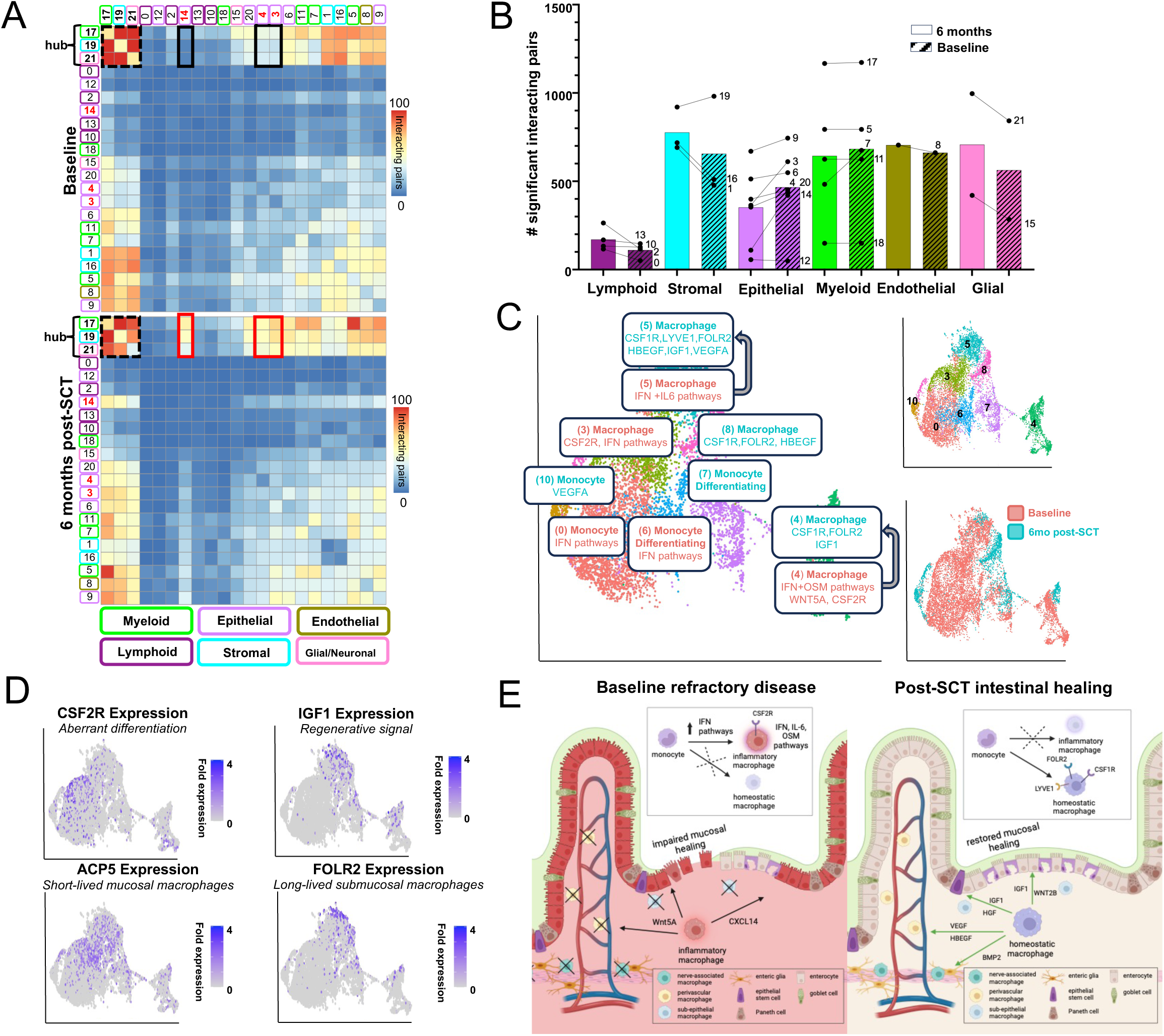
SCT restores homeostatic macrophages in refractory CD. **(A)** scRNA-seq of the intestine was analyzed by CellphoneDB (baseline *n*=4, 6 months post-SCT *n*=3). Significant clusters pairing expression of ligand receptor pairs displayed as a heatmap. Clusters with the largest number of interacting pairs at baseline and 6 months post-SCT are outlined in a dashed black square (i.e. hub clusters). The largest increase in interactions from baseline to 6 months post-SCT are between hub clusters and epithelial clusters 3, 4, 14 outlined in black (baseline) and red (6 months post-SCT). (B) Bar graph (colors per Figure 4A) of mean CellphoneDB interactions per intestinal cell type at baseline and 6 months post-SCT and total interaction for each cluster represented by dot and connecting line. **(C)** UMAP plots from scRNA-seq intestinal dataset with re-clustered myeloid cells annotated (baseline *n*=4, 6 months post-SCT *n*=3). Clusters are highlighted based on abundance at each time point with clusters common to both timepoints that undergo remodeling indicated by arrows (Supplementary Figure 3). Selected significant pathways and marker genes are highlighted for each cluster including differentially expressed genes for clusters common to both timepoints. **(D)** UMAPs demonstrating fold expression of select genes. **(E)** Schematic of monocyte/macrophage populations and functions that reinforce refractory CD pathophysiology at baseline or intestinal healing post-SCT.

When comparing interactions in baseline samples to those obtained 6 months post-SCT the most notable increase occurred between hub clusters 17, 19, and 21 involving epithelial clusters 3 (epithelial stem cell), 4 (enterocyte) and 14 (enterocyte) (Figure 4A). Examination of ligand-receptor pairs revealed signals involving chemokines, extracellular matrix, growth factors and WNT pathway ligands. At baseline, we observed limited interactions predicted to impair epithelial migration and wound healing, while post-transplant we observed large numbers of interactions predicted to support epithelial homeostasis and regeneration. At baseline cellular interactions that impaired epithelial migration, stem cell differentiation and wound healing involved CXCL14/CXCR4 and WNT5A/PTPRK.^33–35^ After SCT there was a complete loss of CXCR4/CXCL14 and WNT5A/PTPRK interactions with a striking increase in interactions involving diverse growth factors including VEGFA, IGF1, HBEGF and HGF that support epithelial stem cell maintenance, epithelial differentiation and epithelial growth.^36–38^ Growth factor signaling extended broadly across myeloid, stromal, glial and epithelial interaction networks post-SCT confirming a specific role for myeloid cells within the immune system to regulate intestinal homeostasis (Supplementary Figure 3).^30, 39–41^

To further understand functional changes among intestinal myeloid cells post-SCT, clusters 5,7,11 and 17 were re-clustered and annotated (Figure 4C). As in the CyTOF analysis there was extensive remodeling post-SCT of intestinal monocyte and macrophage populations suggesting reconstitution of this immune lineage (Figure 4C inset). While certain monocyte/macrophage clusters appeared to be gained or lost post-SCT others were phenotypically remodeled (Figure 4C, Supplementary Figure 3). Pathway analyses and examination of signaling molecules revealed a lineage shift post-SCT from short-lived inflammatory mucosal populations with altered differentiation pathways towards long-lived homeostatic populations with normal differentiation pathways (Figure 4D, Supplementary Figure 3, Supplementary Table 3). Pre-SCT monocyte and macrophage clusters were enriched for interferon signaling pathways with certain macrophage clusters expressing oncostatin M (cluster 4) and IL-6 (cluster 5) signaling pathways associated with medically refractory CD patients.^40, 42, 43^ Cluster 3 and 4 macrophages pre-SCT expressed CSF2RB which promotes aberrant differentiation of inflammatory macrophage populations with impaired restorative functions.^39, 42^ Post-SCT there was significantly increased expression of genes across the macrophage lineage suggesting differentiation of long-lived homeostatic tissue-resident cells expressing restorative signaling molecules (Figure 4C-D). Homeostatic macrophage populations expressed diverse signaling molecules consistent with subepithelial macrophages that maintain epithelial integrity (WNT2B, HGF), perivascular macrophages that support angiogenesis (LYVE1,VEGF) and nerve-associated macrophages that maintain enteric nerves (BMP2) (Figure 4C-E, Supplementary Figure 3).^39, 44^ Among signaling molecules the expression of IGF1 was a marker of macrophage populations enriched post-SCT (Figure 4D). IGF1 and insulin signaling pathways are critical to intestinal healing suggesting absence of these functions in macrophages pre-SCT may impair intestinal healing in medically refractory CD.^45–47^ scRNA-seq analyses confirmed previous studies that suggested differentiation of aberrant macrophage populations reinforce an inflammatory intestinal niche in medically refractory CD.^39, 41^ Our analyses further suggest aberrant macrophage differentiation may lead to loss of restorative functions including growth factor signaling required to coordinate cellular networks necessary for mucosal healing (Figure 4E). As such the therapeutic efficacy of autologous SCT may be due to not only the loss of aberrant inflammatory macrophage populations but the restoration of macrophage populations that support mucosal healing.

### SCT is a myeloid-directed cellular therapy

scRNA-seq and proteomic profiling of immune reconstitution events post-SCT suggested remodeling of intestinal myeloid populations consistent with immune cell reconstitution of this lineage. The ability of SCT to reconstitute (i.e. reset) an immune lineage and function as a cellular therapy requires replacement of resident immune populations from progenitors within the stem cell graft. To characterize early immune reconstitution events post-SCT and interrogate the relationship between immune reconstitution events with progenitor populations we performed scRNA-seq of blood samples at baseline, stem cell collection, engraftment, 3 months post-SCT and 6 months post-SCT.

Cell clusters specific to the time of stem cell collection or engraftment were identified and could not be annotated using gene markers for mature immune cell subsets suggesting that they represent progenitor populations (Figure 5A, Supplementary Figure 4). Three cell clusters (21,24,25) specific to stem cell collection and engraftment expressed CD34 consistent with their being hematopoietic progenitor cells or HSCs (Supplementary Figure 4). To establish ontological relationships between potential progenitor cell clusters and mature immune cell clusters we analyzed scRNA-seq datasets using scVelo (Figure 5A).^48^ scVelo quantifies the ratio of spliced to unspliced RNA (i.e. RNA velocity) to predict ontological relationships between cell populations. RNA velocity analysis confirmed the majority of clusters specific to stem cell collections and engraftment were lineage committed myeloid progenitor populations with relationships to mature circulating myeloid cells. To further understand HSC populations, clusters 21, 24 and 25 were re-clustered and annotated. Re-clustering of CD34^+^ populations distinguished populations specific to stem cell collection (2,4,7), engraftment (0,1,3,6), and common to both timepoints (5,8) (Figure 5B). All CD34^+^ cell clusters were annotated as bone marrow cells confirming the relationship of mobilized and marrow resident HSC populations. CD34^+^ cell clusters included HSC multipotent progenitors (HSC-MPP), cycling HSC-MPP, common myeloid progenitors, megakaryocyte-erythroid progenitors and granulocyte-monocyte progenitors (Figure 5B). Lymphoid-committed CD34^+^ cell clusters were not observed suggesting a bias in mobilized HSCs towards myeloid differentiation.

**Figure 5.**
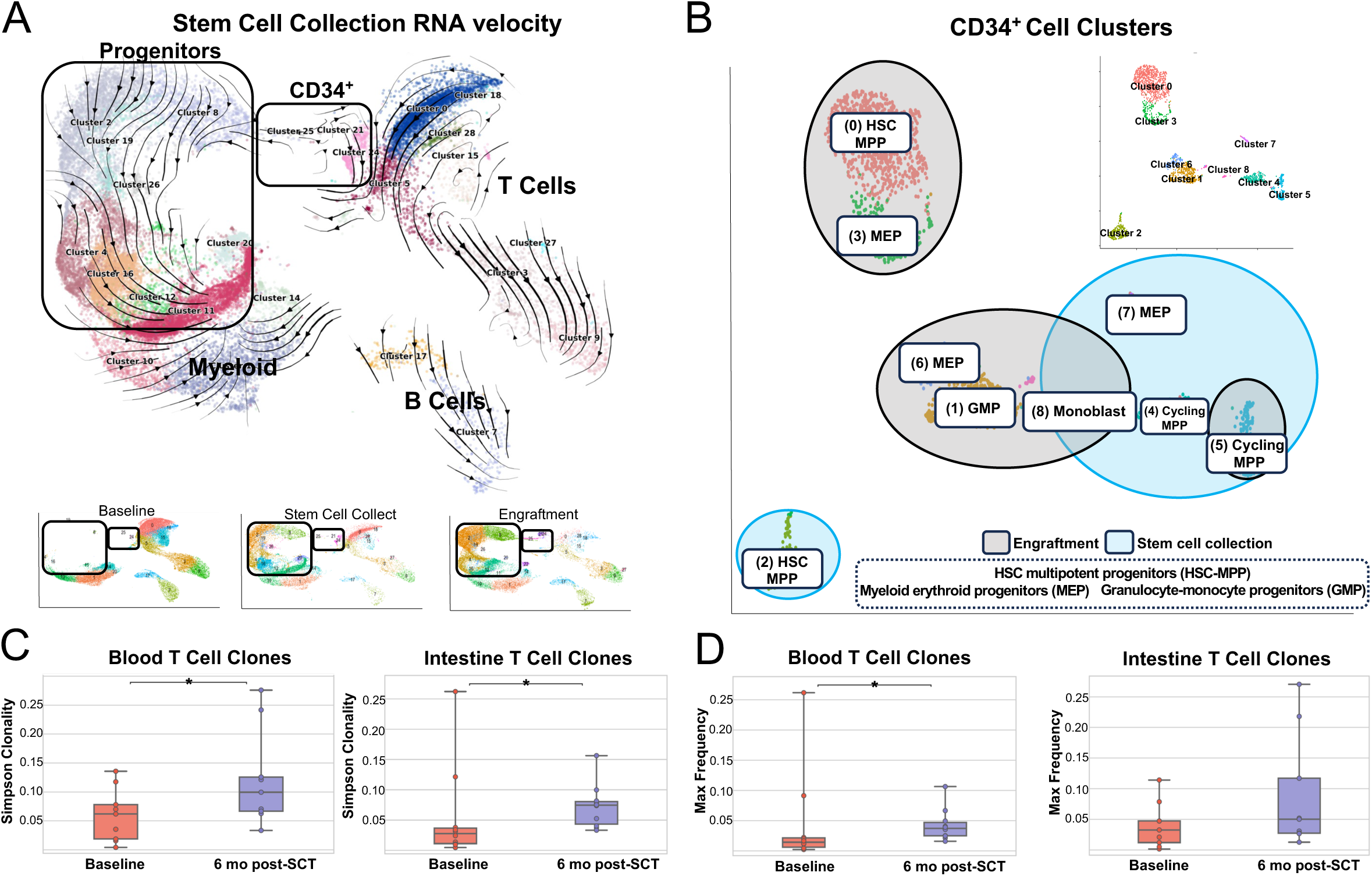
SCT is a myeloid-directed cellular therapy. **(A)** Blood scRNA-seq samples from all timepoints (*n*=20) were clustered with UMAP plots highlighting progenitor clusters specific to stem cell collection (*n*=4) and engraftment (*n*=3) that are not found at baseline (*n*=7). RNA velocity curves using scVelo of stem cell collection demonstrate progenitor clusters relate to mature myeloid cells. **(B)** CD34 expressing clusters were re-clustered and annotated. Clusters are highlighted based on their presence at engraftment or stem cell collection. **(C)** Bar graph (mean ± SEM) of Simpson clonality of TCRβ sequencing of blood (baseline *n*=10, 6 months post-SCT *n*=10) and tissue (baseline *n*=9, 6 months post-SCT *n*=9) samples, Dunn’s test, *P<0.05. **(D)** Bar graph (mean ± SEM) of max clone frequency for T cell clones present at both timepoints, Dunn’s test, *P<0.05.

scRNA-seq analyses suggest that SCT may act as a myeloid-directed cellular therapy driving the early expansion of myeloid progenitors capable of reconstituting intestinal myeloid populations. Analysis of CyTOF datasets support the scRNA-seq analyses and identified lineage negative populations at engraftment that express myeloid markers and integrins necessary for intestinal trafficking (Supplementary Figure 4). To further assess the impact of SCT on lymphoid populations we analyzed changes in T cell clonality through analysis of T cell receptor (TCR) diversity. TCRβ sequencing demonstrated a significant increase post-SCT in TCR clonality in circulating and intestinal populations and a trend towards a decrease in TCR clone diversity post-SCT in the intestine (Figure 5C, Supplementary Figure 5). Analysis of clones present at each timepoint confirmed an increase post-SCT in the frequency of these clones post-SCT suggesting the persistence and expansion of TCR clones post-SCT (Figure 5D). Among the highest frequency clonal populations many clones remained at a high frequency post-SCT especially in the intestine (Supplementary Figure 5). These findings further support that the current SCT protocol may selectively target intestinal myeloid populations and is insufficient to fully reconstitute T cell populations which may be replenished post-SCT through clonal expansion of persistent memory populations and not by progenitors in the stem cell graft.^49, 50^

### HSC heterogeneity in CD

To further interrogate HSC populations and how this may contribute to immune reconstitution post-SCT, we performed functional and phenotypic analyses of the stem cell grafts that were administered to CD patients. Cryopreserved mononuclear cell grafts were obtained from the Mount Sinai Blood Bank and thawed per clinical protocol. Patient samples contained an average of 4.23% CD34^+^ HSCs (range 2.1-8.49%). Phenotypic analysis of CD34^+^ cell populations was performed by flow cytometry to delineate short-term (ST), intermediate-term (IT) and long-term (LT) HSCs and lineage committed progenitors (Figure 6A, Supplementary Figure 6, Supplementary Figure 9). The composition of these CD34^+^ populations varied across patient samples with the predominant populations being ST-HSC or lineage committed progenitors consistent with the scRNA-seq analyses.

**Figure 6.**
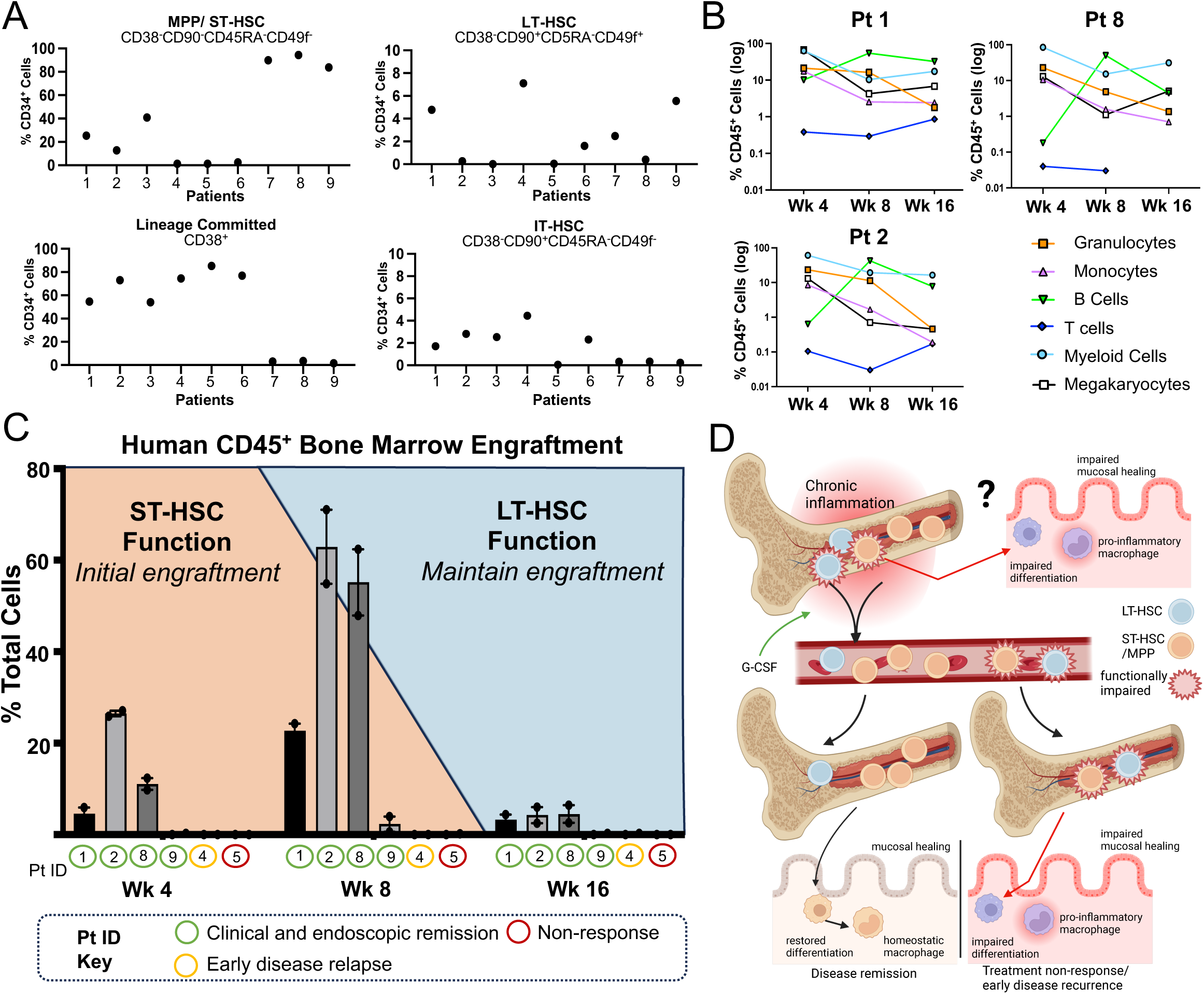
HSC heterogeneity in CD. **(A)** Flow cytometry analysis of mobilized stem cells from 9 patients. Stem cell populations are percent live CD34^+^ cells. **(B)** Human CD34^+^ stem cells were injected into NOD.Cg-*Prkdc^scid^ Il2rg^tm1Wjl^*/SzJ (NSG) mice with mean line graphs of CD45^+^ populations in the bone marrow of the mice from 3 patients with the highest engraftment (Figure 6C) for granulocytes (CD15^+^), monocytes (CD14^+^), B cells (CD19^+^), T cells (CD3^+^), myeloid cells (CD33^+^) and megakaryocytes (CD41a^+^) (mean, *n*=2 biologic replicates at each time point). **(C)** Analysis of engrafted human CD45^+^ cells in the murine bone marrow at 4, 8 and 16 weeks. Bar graph of % human cells from all cells (mean, *n*= 2 biologic replicates at each time point). Pt ID key identifies clinical response for each patient. Graph background color demonstrates the contribution of ST-HSC and LT-HSC to human engraftment at each timepoint. **(D)** Schematic of hypothesis that chronic inflammation may impair HSC functions perpetuating refractory disease and affecting SCT outcomes.

To functionally interrogate HSC grafts that were administered to the CD patients, their CD34^+^ cells were transplanted into NOD.Cg-*Prkdc^scid^ Il2rg^tm1Wjl^*/SzJ (NSG) mice.^51^ HSC engraftment and cell differentiation potential was assessed at week 4, 8 and 16 within the bone marrow, spleen, and peripheral blood of recipient mice using flow cytometry (Figure 6B-C, Supplementary Figure 6, Supplementary Figure 9). Examination of the human cell subpopulations confirmed the presence of functional HSCs capable of multilineage hematopoiesis (Figure 6B, Supplementary Figure 6). Analysis of mature CD45^+^CD34^-^ cells at week 4 indicated that CD33^+^ myeloid cells were the predominant cell population in the bone marrow. At week 8 there was an increase in CD19^+^ cells which decreased by week 16 with CD33^+^ cells again being the dominant population. Mature CD3^+^ T cells were detected only transiently and at exceedingly low levels in all mice including in the spleen and blood (Supplementary Figure 6).^52^ The dynamics of immune reconstitution events in the xenograft model paralleled patient observations where myeloid progenitors emerged at engraftment followed by a rise in naïve B cell populations and a persistent decrease in naïve T cell populations (Supplementary Figure 2).

At each time point, the engraftment of human CD45^+^ cells in the bone marrow of recipient mice ranged from 0-71% (Figure 6C). The greatest degree of human engraftment was observed at week 8 and reduced by week 16. Engraftment of human CD34^+^ cells in the bone marrow was similarly reduced by week 16 likely reflecting the limited number of functional LT-HSCs present in these grafts (Supplementary Figure 6).^53^ The observed functional heterogeneity of HSC populations in the xenograft model despite classification using established protein markers suggests dysfunction in HSC differentiation and repopulation capacity especially among LT-HSC.^52, 54^ Correlation between HSC function in the xenograft model and SCT clinical outcomes is not feasible due to the rarity of clinical non-response in this study at 6 months (1/14 patients). In follow up to 12 months a second patient had a clinical/endoscopic relapse 8 months post-SCT. It was observed that stem cell grafts from the two patients with a negative SCT clinical outcome failed to engraft in the xenograft model whereas stem cell grafts from four patients with stable endoscopic and clinical outcomes post-SCT achieved human cell chimerism in the xenograft model (Figure 6C). The functional heterogeneity of HSC from CD patients in the xenograft model is inconsistent with historical observations in healthy patients suggesting a systemic impact of CD on HSC populations.^52–54^

## Discussion

Our observations suggest that SCT functions as a myeloid-directed cellular therapy in CD involving HSC mediated reconstitution of intestinal macrophages capable of supporting mucosal healing. These observations are independently validated using multimodal studies of immune reconstitution events including CyTOF and scRNA-seq. Functional interrogation of HSCs in xenograft models further supports that HSCs in SCT protocols shape clinical outcomes including the timing of immune reconstitution, the selected reconstitution of specific cell lineages and potentially the clinical efficacy of SCT. The observations in the xenograft models that graft mediated reconstitution of the marrow niche may be a critical therapeutic outcome suggests the inability to repopulate HSC populations in the marrow may underlie the consistent clinical observation that SCT is incapable of maintaining long-term disease free remission in many patients.^4, 5, 55–57^ Future studies in larger cohorts over longer periods of time that capture events related to treatment non-response and CD recurrence will be critical to validate the therapeutic mechanisms proposed in this study and whether HSC heterogeneity impacts SCT outcomes. The heterogeneity of HSC populations in CD as determined by functional analyses of the stem cell grafts further suggests CD pathophysiology may create a dysfunctional marrow niche shaped by the CD inflammatory milieu. The functional impairment in xenograft models of ST-HSC and LT-HSC populations is consistent with studies that demonstrate an impact of chronic inflammation on these HSC populations.^55, 58, 59^ The importance of how CD affects the function of specific HSC populations likely extends beyond therapeutic implications proposed here and may be critical to how HSC populations reinforce disease pathophysiology especially in the myeloid lineage. It is interesting to consider that aberrant intestinal macrophage populations in medically refractory CD may be perpetuated through acquired changes in differentiation programs among marrow progenitor populations necessitating cellular therapies capable of targeting these populations (Figure 6D).^56, 57, 60, 61^

The successful use of SCT as a cellular therapy requires an understanding of how hematopoietic progenitors shape tissue resident immune populations. The intestinal myeloid niche is largely dependent on circulating progenitor populations and as such may be more amenable to reconstitution with SCT.^27, 39^ The inability of the current SCT protocol to adequately reconstitute the intestinal lymphoid lineages, especially T cells, may be secondary to the observed myeloid bias of the harvested stem cell grafts, age related changes in hematopoiesis and limitations of the conditioning chemotherapy to deplete tissue resident lymphoid memory cell populations.^49, 50, 61, 62^ It is possible the persistence of pathogenic intestinal T cell clones post-SCT may impact long-term clinical outcomes which will require further study. While minimal changes among intestinal lymphoid populations were observed, significant changes did occur in circulating lymphoid populations, especially B cells. These findings reinforce the importance of profiling immune reconstitution events across disease specific immune niches and suggest the efficacy of SCT for B cell mediated diseases including multiple sclerosis and systemic sclerosis may derive from reconstitution of the circulating B cell compartment.^63, 64^ Patients in this study were exposed to vedolizumab after immune reconstitution requiring additional studies to deconvolute the effect of vedolizumab from our observations. A previous study of SCT in CD patients who were not exposed to vedolizumab reported similar changes in the circulating B cell compartment.^8^ With a greater understanding of the interplay between SCT conditioning and mobilization regimens with HSC mediated immune reconstitution events, we anticipate next generation SCT protocols will safely and efficiently target pathogenic immune cell lineages and sustain immune reconstitution events over time.

The last two decades have brought an unprecedented growth in CD therapies, but it has remained consistent that each new therapy is less effective for patients that have failed a previous therapy.^14, 65, 66^ While CD therapies seemingly encompass distinct therapeutic mechanisms, all therapies target the inflammatory response supporting the concept that medically refractory disease involves pathways not directly related to inflammation. This study suggests SCT acts as myeloid-directed cellular therapy restoring macrophage functions critical to intestinal healing. That the pathophysiology of medically refractory disease involves defects in cellular pathways required to heal the intestine after resolution of inflammation provides a plausible hypothesis for why even novel immunotherapies fail in medically refractory patients.^16, 67^ The findings in this study further reinforce the central role of intestinal macrophages in CD biology including medically refractory disease.^39–42^ An additional factor that might contribute to the recurrence of CD after SCT is the inability of autologous HSCs to correct genetically driven defects in intestinal macrophages.^68^ The studies presented here suggest the genetic correction of HSC in SCT grafts may be sufficient to restore wild-type function of resident myeloid cells in the intestine of CD patients leading to long-term disease free remission.^69^ While the widespread use of SCT in CD patients is limited by toxicity, this study adds to the growing literature confirming the clinical efficacy of SCT for patients with medically refractory CD. The therapeutic applications of observations made in this study of SCT may be generalizable to all patients with CD through the future development of regenerative cellular or small molecule/biologic therapies that support tissue healing and not only suppress the inflammatory response.^70^

## ABBREVIATIONS

ATG: thymoglobulin
CDAI: Crohn’s Disease Activity Index
FDR: false discovery rate
HBI: Harvey Bradshaw Index
HSC: hematopoietic stem cell
IT-HSC: intermediate-term hematopoietic stem cell
LT-HSC: long-term hematopoietic stem cell
MPP: multipotent progenitor
NSG: NOD.Cg-Prkdc^scid^ *Il2rg^tm1Wjl^*/SzJ
SAM: significance of microarray
ScRNA-seq: single cell RNA sequencing
SES-CD: Simplified Endoscopic Score for Crohn’s Disease
SCT: Stem cell transplant
ST-HSC: short-term hematopoietic stem cell
TCR: T-cell receptor
UMAP: uniform manifold projection

## AUTHOR CONTRIBUTIONS

Conceptualization – L.J.C.

Methodology – L.J.C, J-F.C., J.C, R.H.

Software – S.T., C.T.

Validation – all authors

Formal Analysis – D.G., S.T., X.W., L.J.C.

Investigation – D.G., S.T., X.W., A.D., A.E., E.F., L-S.C., K.S., S.S.

Resources – X.W., A.D., A.E., J-F. C., C.T., J.E.L., J.L.M.F., R.H., J.C., L.J.C.

Data Curation – D.G., S.T., E.F., S.S., L.J.C.

Writing – Original Draft – D.G., L.J.C

Writing – Review & Editing, all authors Visualization – D.G., L.J.C.

Supervision, A.E., J-F. C., C.S., J.E.L., J.L.M.F., B.K.M., R.H., J.C., L.J.C

Project Administration, D.G., L.J.C. Funding Acquisition, L.J.C.

## ACKNOWLEDGEMENTS

At the Icahn School of Medicine we acknowledge the Human Immune Monitoring Center for assistance in the CyTOF assays. This work was supported in part through the computational and data resources and staff expertise provided by Scientific Computing and Data at the Icahn School of Medicine at Mount Sinai and supported by the Clinical and Translational Science Awards (CTSA) grant UL1TR004419 from the National Center for Advancing Translational Sciences. This study was funded by the Leona M. and Harry B. Helmsley Charitable Trust (L.J.C.). We would like to acknowledge Dr. Camelia Iancu-Rubin and Dr. Suzanne A. Arinsburg from the Cellular Therapy Laboratory at Mount Sinai for their assistance in acquiring and processing peripheral mobilized blood stem cell collections (IRB protocol STUDY-21-0062). We would like to acknowledge the MASCT CD study team; Alema Gonzalez, Ladislao Decenteceo, Jonathan Lagdameo, Moutasem Mansi, Stephanie Gold, David Faleck, Amir Steinberg, Saurabh Mehandru, Bruce Sands, Ryan Ungaro, Alexander Greenstein, Laurie Keefer, and Laura Manning. We thank the Duke University School of Medicine for the use of the Sequencing and Genomic Technologies Shared Resource, which provided sequencing service for scRNA-seq samples. All graphical illustrations were created with BioRender.com

## Data Transparency Statement

### Materials availability

This study did not generate new unique reagents

### Data and code availability

- Raw and processed scRNA-seq data will be deposited
- Data reported will be shared by the lead contact upon request
- No new code was generated. All R scripts are available from the lead contact upon request
- Any additional information required to reanalyze the data reported is available from the lead contact upon request

