## Supplementary Methods for "The reparative immunologic consequences of stem cell transplantation as a cellular therapy for refractory Crohn’s disease"

**SUPPLEMENTARY MATERIALS**

**Supplementary Figure 1**

**CyTOF supervised.**

**(A)** Supervised clustering of immune cell populations from intestinal (baseline, *n*=13, 6 months post-SCT *n*=13) and blood samples (baseline *n*=9, 6 months post-SCT *n*=11) as mean percentage of total cells. Heat map demonstrates the log ratio of change at 6 months post-SCT compared to baseline for intestine (top) and blood (bottom). **(B)** Bar graph (mean ± SEM) of selected FlowSOM metaclusters at baseline and 6 months post-SCT in intestine (baseline *n* =13, 6 months post-SCT *n* =13) and blood (baseline, *n* =7, 6 months post-SCT *n* =9); Mann-Whitney test, ** P<0.01.

**Supplementary Figure 2**

**CyTOF intestinal changes.**

**(A)** viSNE analysis of manually gated populations from intestinal samples (baseline *n*=13, 6 months post-SCT *n*=13). All cells are concatenated as a single file. **(B)** The dynamics of manually gated immune populations over time. Line graph expressed as log fold change relative to baseline (mean ± SEM; baseline *n*=9, stem cell collection *n*=5, engraftment *n*=3, 3 months post-SCT *n*=6, 6 months post-SCT *n*=12, 12 months post-SCT *n*=11).

**Supplementary Figure 3**

**scRNA-seq of intestine.**

**(A)** UMAP plot of intestine samples combined baseline (*n*=4) and 6 months post-SCT (*n*=3). **(B)** Dotplot of growth factors expressed in scRNA-seq dataset highlighting expression in myeloid clusters and lymphoid clusters. Percent expressing cells correlates with size of dot and color correlates with average expression to heatmap. **(C)** UMAP plot of re-clustered intestinal myeloid cells. **(D)** UMAP plot colored by samples from baseline and 6 months post-SCT. **(E)** Table of myeloid clusters with mean population frequencies at baseline and 6 months post-SCT. Clusters enriched at specific time points are highlighted. **(F)** Dotplot of growth factor, growth factor receptors and WNT ligands expressed in the CellphoneDB database in myeloid clusters with select ligands/receptors from Figure 4E highlighted by a dashed box.

**Supplementary Figure 4**

**scRNA-seq of blood.**

**(A)** Combined UMAP plot of blood samples from all timepoints (*n*=20) (left) and separated by baseline (*n*=7), stem cell collection (*n*=4) and engraftment (*n*=3) (right). **(B)** scRNA-seq clusters from Extended Data Fig. 4a as percentage of total cells at each time point. **(C)** scRNA-seq UMAP of samples from blood at stem cell collection and engraftment demonstrating CD34 expression (*n*=4 stem cell collection, *n*=3 engraftment). **(D)** viSNE analysis of live cells from blood samples using CyTOF taken at baseline and engraftment. All cells are concatenated as a single file (*n*=7 baseline, *n*=3 engraftment). Individual markers shown as heatmaps for viSNE analysis at time of engraftment.

**Supplementary Figure 5**

**TCRβ sequencing.**

**(A)** Bar graph (mean ± SEM) of Simpson diversity of TCRβ sequencing of blood (baseline *n*=10, 6 mo post-SCT *n*=10) and tissue (baseline *n*=9, 6 mo post-SCT *n*=9) samples at baseline and 6 mo post-SCT, Wilcoxon matched paired t-test, ns P>0.05, * P< 0.05, ** P<0.01. Higher Shannon diversity value represent a decrease in clone diversity. **(B)** Bar graph (mean ± SEM) of the number of clones present in the top 10 high frequency clones at both baseline and 6 months post-SCT for each patient in the blood and the tissue.

**Supplementary Figure 6**

**Stem cell graft and xenograft model.**

**(A)** Flow cytometry analysis of mobilized stem cells from 9 patients. Stem cell populations are percent live CD34^+^ cells. The most abundant CD34^+^ cell population for each patient are highlighted in green. Human CD34^+^ stem cells were injected into NOD.Cg-*Prkdc^scid^ Il2rg^tm1Wjl^*/SzJ (NSG) mice with analysis of engrafted human cells in the bone marrow **(B),** spleen **(C)**, and peripheral blood **(D)** at 4, 8 and 16 weeks. Mean bar graph of % human cells among all cells for each body site (*n*= 2 biologic replicates at each time point for each patient sample). Pt ID key identifies clinical response for each patient. **(E)** Mean line graphs for % of each mature populations from human CD45^+^ cells in the spleen (top) and peripheral blood (bottom) of the mice from 3 patients with engraftment (>0.1%) for granulocytes (CD15^+^), monocytes (CD14^+^), B cells (CD19^+^), T cells (CD3^+^), myeloid cells (CD33+) and megakaryocytes (CD41a+) (mean, log scale).

**Supplementary Figure 7**

**Blood CyTOF - CITRUS tree markers**

**Supplementary Figure 8**

**Intestine CyTOF - CITRUS tree markers**

**Supplementary Figure 9**

**Flow cytometry gating for stem cell graft and xenograft model**

**(A)** Flow cytometry gating strategy of patient stem cell graft and humanized cells from xenograft mouse model for samples from the bone marrow, blood and spleen to resolve stem cell populations and **(B)** mature immune cell populations.

**Supplementary Table 1: CyTOF cluster medians**

**Supplementary Table 2: CellphoneDB interactions**

**Supplementary Table 3: scRNA-seq WikiPathways analysis**

**Supplementary Table 4: Bulk RNA-seq WikiPathways analysis**

**Supplementary Methods**

MASCT-CD Trial

Adult patients >18 years old with a diagnosis of CD and active clinical (Crohn’s Disease Activity Index (CDAI) > 250) and endoscopic disease (Simple Endoscopic Score for Crohn’s Disease (SES-CD) >3 in at least 1 segment) were enrolled.^1^ Inclusion criteria included failure to respond to a member of each of the following drug classes: corticosteroids, immunomodulators (azathioprine, 6-mercaptopurine, methotrexate), anti-TNFα, vedolizumab and ustekinumab and without any reasonable surgical option. Each patient was reviewed by a steering committee comprised of IBD gastroenterologists, surgeons and BMT physicians to ensure each patient met inclusion criteria, the SCT would likely provide potential benefit and there were no comorbid conditions that put the participant at excessive risk. Participants were required to discontinue immunosuppressive medication prior to mobilization based on standard washout periods or demonstrated drug level of 0.^2^ Prednisone was discontinued or tapered to physiologic replacement dose of 5-10mg/day. Enrolled participants received stem cell mobilization with cyclophosphamide 2 g/m^2^/day for 2 days followed by G-CSF 10 mg/kg/day with daily leukapheresis until collection goal of >20x10^4^/ml CD34^+^ cells. The conditioning regimen included cyclophosphamide 50 mg/kg/day from day -6 to day -3 and then ATG (thymoglobulin) 2.5 mg/kg/day from day -3 to day -1 prior to SCT. After stem cell engraftment (neutrophil count ≥0.5x10^3^/μL for 3 days) participants received vedolizumab 300mg within 3 days of discharge with standard induction and maintenance doses.^3^ Participants were evaluated at baseline, stem cell collection, stem cell engraftment and longitudinally until end of study at 54 weeks. Samples of blood, intestinal biopsies, and clinical symptoms (ie Harvey Bradshaw Index (HBI)) were collected longitudinally throughout.^4, 5^

PBMC isolation

Blood was collected into CPT tubes (BD Vacutainer) and cells were isolated within 2 hours of blood draw. The tubes were centrifuged for 20 min at 1800G at room temperature without break and then the buffy coat layer of PBMCs was collected and then washed with PBS, centrifuged for 15 minutes at 300G. The pellet was resuspended in sterile PBS and viability and cell count were determined using AO/PI dye and Nexcelcom automated cell counter prior to further processing for downstream analyses. Sample viability was routinely >90%.

Intestinal lamina propria single cell isolation

Tissue biopsies were taken endoscopically with biopsy forceps and collected in ice cold complete RPMI 1640 supplemented with glutamine, 1x pen/strep, 10mM HEPES and 10% FBS and processed within 1 hour of collection. The biopsy samples were pooled together, 10-20 biopsies per sample. Epithelial cells were dissociated by transfer into dissociation medium (HBSS w/o Ca^2+^ Mg^2+^, with HEPES 10mM, EDTA 5mM) and agitated for 30 minutes at 37°C at 100rpm, and then vortexed for 30 seconds and then filtered through 70mm strainer. The biopsies were then washed with complete RMPI 1640 at room temperature and then transferred to digestion media (HBSS with Ca^2+^ Mg^2+^, FCS 2%, DNase I 0.5mg/ml,Collagenase IV 0.5mg/ml) and agitated for 40 minutes at 37°C at 100rpm. After digestion the suspension was filtered through a 70mm strainer and washed with complete RPMI, spun down 5 minutes at 2500rpm and resuspended in 1ml red blood cell lysis buffer and incubated for 10 minutes at room temperature, spun again for 5 minutes at 2500 rpm and then resuspended in RPMI and viability and cell count were determined using AO/PI dye and Nexcelcom automated cell counter prior to further processing for downstream analyses. For samples with viability 70% or less, dead cells were depleted using a dead cell removal kit (Miltenyi Biotec) following the protocol from the manufacturer. After dead cell removal, viability was routinely >85%.

CyTOF sample preparation and analysis

Isolated cells were washed with FACS buffer and fixed with 1.6% paraformaldehyde at room temperature. Cells were then washed twice in FACS buffer and placed at 4°C until staining and analysis by CyTOF. Prior to CyTOF assays previously optimized antibody mixtures were prepared in cell staining media (CSM, Fluidigm). Antibodies are outlined in key resources table. Each sample was washed and resuspended in 800 μl of 1X Barcode Perm Buffer (Fluidigm). Compatible Pd-barcodes were thawed, resuspended in 100 μL of 1X Barcode Perm Buffer and added to the samples. Samples were incubated on ice for 30 minutes then washed in CSM and pooled together. Each barcoded set of samples was resuspended in 100 μL of CSM containing 100 U/ml heparin (Sigma) to block non-specific MaxPar Antibody binding. A titrated surface antibody panel designed to allow identification of all major immune subsets was prepared in an additional 100 μl of CSM, filtered through a 0.1 μm spin filter (Amicon) and added directly to the sample. Samples were stained for 30 minutes on ice, then washed with CSM and fixed with freshly diluted 2% formaldehyde (Electron Microscopy Sciences) in PBS to cross-link and preserve all surface antibodies. The samples were washed and stored as pellets in CSM until CyTOF acquisition. Immediately prior to acquisition, samples were washed once with PBS, once with deionized water and then counted and resuspended at a concentration of 1 million cells/ml in water containing a 1/20 dilution of EQ. 4 Element Beads (Fluidigm).

CyTOF2 Mass Cytometer equipped with a SuperSample fluidics system (Victorian Airships) to facilitate bulk sample acquisitions. Samples were acquired at a flow rate of 0.045 mL/min and an event rate of < 400 events per second.

For CyTOF analysis cell events were identified as Ir191/193 positive events, and residual Ce140^+^ normalization beads were excluded.

Supervised analyses used the following number of samples: blood (n= 9 baseline, n=5 stem cell collection, n=4 engraftment, n=5 at 3 months post-SCT, n=11 at 6 months post-SCT and n=7 at 12 months post-SCT), tissue (n=13 baseline, n=13 at 6 months post-SCT). Supervised analysis was performed using protein markers shared between circulating and intestinal immune panels to facilitate comparison and used previously reported gating strategy.^6^

CyTOF unsupervised analyses (viSNE, FlowSOM, CITRUS) used the following number of samples: tissue analysis (n=13 baseline, n=13 at 6 months post-SCT), Blood analysis (n=9 baseline, n=4 at engraftment, n=11 at 6 months follow up). FlowSOM analyses were performed with 50,000 events per sample clustering with all channels using hierarchical consensus methods. After iterative analyses 15 metaclusters and 100 clusters were chosen for unsupervised clustering based on correlation with canonical markers suggesting resolution of meaningful populations. CITRUS analysis was performed using all channels with equal event sampling, 5000 events per file and minimum cluster size of 1%. The significance of microarray (SAM) model was utilized with a false discovery rate (FDR) of 0.01. viSNE analysis was performed on concatenated samples. Principal Component Analysis (PCA) was applied to FlowSOM generated cell clusters and we generated PCA plots using R and the ggplot2 package with the create_pca_plot function. Ellipses representing a 95% confidence interval were used.

Flow Cytometry of stem cell graft

Patient mobilized mononuclear cells were obtained from the Laboratory of Cellular Therapy at Mount Sinai Hospital. Cells were thawed per clinical protocol in a 37°C water bath and washed with PBS with 2% FBS solution, centrifuged for 5 minutes at 300G and then incubated with RBC lysis buffer (G-Biosciences) for 5 minutes at room temperature, centrifuged for 5 minutes at 300G followed by another wash and subsequently processed for analysis by flow cytometry using the Attune NxT Flow Cytometer (ThermoFisher). List of antibodies is provided in the key reseource table and gating strategy is provided in Supplementary Figure 9.

Xenograft

All mice were housed in environmentally controlled animal facilities with unrestricted access to food and water.

Peripheral mobilized stem cells underwent CD34+ magnetic bead selection with the Easy Sep Human CD34 Selection Kit II (StemCell Technologies) and live CD34+ purity was confirmed via flow cytometry and was routinely ≥90%.

To determine the presence of human cells belonging to various hematopoietic lineages in the recipient mice, cells from were stained with a panel of antibodies specific to human antigens (key resources table) and analyzed using the Attune NxT Flow Cytometer (ThermoFisher Scientific). Cells obtained from mice not receiving transplants were analyzed in a similar fashion to exclude the possibility of cross-reactivity of the monoclonal antibodies with murine cells.

Single Cell RNA sequencing sample processing

After single cell isolations were obtained the cells were made into suspension with 10,000 cells/sample loaded onto the Chromium Next GEM Chip G (10x Genomics, Chromium Single Cell 3’ v3.1 kit) with GemCode Gel Beads and run on the 10x Genomics Chromium iX instrument and samples were processed per manufacturers protocol. Libraries were prepared using 10x Genomics Library Construction Kit and QC was completed with the Agilent Bioanalyzer High Sensitivity DNA kit prior to submitting for sequencing on the Illumina NovaSeq 6000 at the Duke Center for Genomic and Computational Biology at Duke University and they provided FASTQ files.

Single cell data processing

Seurat object was created using the gene expression (GEX) modality with implementing filters for a minimum of 3 cells and 200 features per barcode.^7^ Object was refined with parameters such as 500 < max.count.rna < 60000, max.features < 2000, and max.percent.mt < 30.

To stabilize gene expression variance, the SCTransform v2 function was applied, integrating normalization, variable feature selection, and data scaling into a single operation. The normalized GEX outputs underwent PCA and uniform manifold projection (UMAP) for dimensionality reduction and 2-dimensional projection, utilizing 20 dimensions. Neighborhood formation and clustering were performed using 20 dimensions at a resolution of 0.5.

To address batch effects, individual samples were consolidated and harmonized into a unified object using the Harmony package. The distinct Seurat objects were merged using the base R merge function and subsequently subjected to SCTransform and PCA reduction. The RunHarmony function was then applied to the reduced output, grouping the data by individual objects. Following harmonization, the manifold projection, neighborhood formation, and clustering steps were repeated using the unchanged parameters.

In Seurat the Findallmarkers function was used to find positively differentially expressed genes for each cluster that were significantly different (P<0.05) from all other clusters. This function was used to find marker genes and to compare clusters at different time points. Dot plots were created using the DotPlot function using the Seurat object for selected genes of interest where the color represents the average expression and the size represents the percentage of cells expressing that gene.

RNA velocity^8^

Python based implementation of the Velocyto (v0.17) was used to generate loom files, capturing the splicing kinetics of genes. The loom files were integrated into Seurat objects, ensuring compatibility for downstream analysis using scVelo (v0.2.4) to model transcription dynamics (stochastic model) and velocities were visualized using stream plots.

CellphoneDB^9^

Ligand-receptor analysis was conducted using CellPhoneDB v2.0 with the statistical_analysis function using the counts matrix and metadata from the generated Seurat objects. Heatmaps were generated using ktplots, a set of R plotting functions designed for visualizing gene expression data in single-cell datasets.

Single Cell RNA sequencing annotation

Single cell UMAP clusters were annotated using known sets of marker-genes to identify major cell types using previously annotated reference data sets including Gut Cell Atlas, CZ GENExCELL atlas and Enrichr cell types with reference data bases PanglaoDB, CellMarker and Human Gene Atlas.^7, 10-13^. Additionally, HSC and hematopoietic progenitor populations were annotated using the CZ CELLxGENE bone marrow dataset as a reference.^14, 15^

TCR sequencing

Isolated PBMCs and intestinal biopsy samples that were collected and stored in RNAlater (Sigma Aldrich) were sent to Adaptive Biotechnologies for further processing which included their in-house protocol for genomic DNA extraction followed by TCRβ sequencing with survey resolution for tissue samples (≤10^5^ T cells) and deep resolution for PBMC samples (>10^5^ T cells). The somatically rearranged CDR3 region was amplified from genomic DNA using a two-step, amplification bias-controlled multiplex PCR approach^16^. CDR3 libraries were sequenced on an Illumina sequencing instrument. Data from Adpative Biotechnologies was analyzed and extracted using the provided immunoSEQ analyzer.

**Key resources table**

| REAGENT or RESOURCE | SOURCE | IDENTIFIER |
| --- | --- | --- |
| **Antibodies** |  |  |
| APC anti-human CD34 antibody (clone 581) | BD | Cat#555824 |
| FITC anti-human CD38 antibody (clone HIT2) | BD | Cat#555459 |
| super bright 436 anti-human CD90 antibody (clone eBio5E10 (5E10)) | Invitrogen | Cat#62-0909-42 |
| APC/Cyanine7 anti-human CD49f antibody (clone GoH3) | BioLegend | Cat#313627 |
| eFluor™ 506 anti-human CD45RA (clone HI100) | Invitrogen | Cat#69-0458-42 |
| Alexa Fluor™ 700 anti-human CD33 (clone WM-53) | Invitrogen | Cat#56-0338-42 |
| PE anti-human CD201 (clone RCR-401) | BioLegend | Cat#351904 |
| FITC anti-human CD45 (clone HI30) | BD | Cat#BDB555482 |
| PE anti-human CD135 (clone 4G8) | BD | Cat#BDB558996 |
| PerCP-Cy5.5 anti-human CD33 (clone P67.6) | BD | Cat#BDB341650 |
| PE-Cyanine7 anti-human CD34 (clone 4H11) | Invitrogen | Cat#5015502 |
| APC anti-human CD49f (clone GoH3) | BioLegend | Cat#50166770 |
| Alexa Fluor™ 700 anti-human CD7 (clone 124-1D1) | Invitrogen | Cat#5016858 |
| APC/Cyanine7 anti-human CD38 (clone HIT2) | BioLegend | Cat#50403232 |
| Super Bright™ 436 anti-human CD90 (clone 5E10) | Invitrogen | Cat#62090942 |
| Super Bright™ 600 anti-human CD45RA (clone HI100) | Invitrogen | Cat#63045842 |
| Super Bright™ 702 anti-human CD10 (clone CB-CALLA) | Invitrogen | Cat#67010642 |
| FITC anti-human CD19 (clone HIB19) | Invitrogen | Cat#509476 |
| PE anti-human CD33 (clone WM53) | BD | Cat#BDB555450 |
| PerCP-eFluor™ 710 anti-human SSEA1 (clone MC-480) | Invitrogen | Cat#46881342 |
| APC anti-human CD45 (clone HI30) | Invitrogen | Cat#5014986 |
| Alexa Fluor™ 700 anti-human CD14 (clone 61D3) | Invitrogen | Cat#501124675 |
| Super Bright™ 436 anti-human CD41a (clone HIP8) | Invitrogen | Cat#62041942 |
| Super Bright™ 600 anti-human CD3 (clone OKT3) | Invitrogen | Cat#63003742 |
| Super Bright™ 702 anti-human CD235a (clone HIR2 (GA-R2)) | Invitrogen | Cat#67998742 |
| anti-human CD45-89Y (clone HI30) | Fluidigm | Cat#3089003B |
| anti-human CD57-113In (clone HNK-1) | Biolegend | Cat#359602 |
| anti-human CD11c-115In (clone REA618) | Miltenyi | Cat#130-122-296 |
| anti-human CD326-115In (clone 9C4) | Fluidigm | Cat#3141006B |
| anti-human CD19-142Nd (clone HIB19) | Fluidigm | Cat#3142001B |
| anti-human CD45RA-143Nd (clone HI100) | Fluidigm | Cat#3143006B |
| anti-human CD141-144Nd (clone Phx-01) | Biolegend | Cat#902101 |
| anti-human CD4-145Nd (clone REA623) | Miltenyi | Cat#130-122-283 |
| anti-human CD8-146Nd (clone RPA-T8) | Fluidigm | Cat#3146001B |
| anti-human IgA-147Sm (polyclonal) | SouthernBiotech | Cat#2050-01 |
| anti-human CD16-148Nd (clone 3G8) | Fluidigm | Cat#3148004B |
| anti-human CD127-149Sm (clone A019D5) | Fluidigm | Cat#3149011B |
| anti-human CD1c-150Nd (clone L161) | Biolegend | Cat#331502 |
| anti-human CD123-151Eu (clone REA918) | Miltenyi | Cat#130-122-297 |
| anti-human CD66b-152Sm (clone REA306) | Miltenyi | Cat#130-108-019 |
| anti-human PD-1-153Eu (clone EH12.2H7) | Biolegend | Cat#329926 |
| anti-human CD86-154Sm (clone IT2.2) | Biolegend | Cat#305449 |
| anti-human CD27-155Gd (clone REA499) | Miltenyi | Cat#130-122-295 |
| anti-human CXCR3-156Gd (clone G025H7) | Fluidigm | Cat#3156004B |
| anti-human CD33-158Gd (clone WM53) | Fluidigm | Cat#3158001B |
| anti-human CD103-159Tb (clone Ber-ACT8) | BioLegend | Cat#350202 |
| anti-human CD14-160Gd (clone REA599) | Miltenyi | Cat#130-122-290 |
| anti-human CD56-161Dy (clone REA196) | Miltenyi | Cat#130-108-016 |
| anti-human CD64-162Dy (clone 10.1) | Biolegend | Cat#305002 |
| anti-human CD172ab-163Dy (clone SE5A5) | Fluidigm | Cat#3163017B |
| anti-human CD69-164Dy (clone FN50) | Biolegend | Cat#310902 |
| anti-human FceRIa-165Ho (clone AER-37) | BioLegend | Cat#334602 |
| anti-human CD25-166Er (clone REA570) | Miltenyi | Cat#130-122-302 |
| anti-human CD3-168Er (clone REA613) | Miltenyi | Cat#130-122-282 |
| anti-human Beta7-169Tm (clone REA441) | Miltenyi | Cat#130-122-309 |
| anti-human CD38-170Er (clone REA671) | Miltenyi | Cat#130-122-288 |
| anti-human CD161-171Yb (clone HP-3G10) | Biolegend | Cat#339902 |
| anti-human CD206-172Yb (clone 15-2) | Biolegend | Cat#321102 |
| anti-human CXCR4-173Yb (clone 12G5) | Fluidigm | Cat#3173001B |
| anti-human HLADR-174Yb (clone REA805) | Miltenyi | Cat#130-122-299 |
| anti-human PD-L1-175Lu (clone 29E.2A3) | Fluidigm | Cat#3175017B |
| anti-human CD54-176Yb (clone HCD54) | BioLegend | Cat#353102 |
| anti-human CD11b-209Bi (clone ICRF44) | Fluidigm | Cat#3209003B |
| anti-human CD57-113In (clone HNK-1) | BioLegend | Cat#359602 |
| anti-human CD45-115In (clone HI30) | BioLegend | Cat#304002 |
| anti-human Va7.2-141Pr (clone 3C10) | BioLegend | Cat#351702 |
| anti-human CD62L-144Nd (clone DREG-56) | BioLegend | Cat#304802 |
| anti-human CD49d-147Sm (clone 9F10) | BioLegend | Cat#304310 |
| anti-human BTLA-158Gd (clone MIH26) | BioLegend | Cat#344502 |
| anti-human CCR4-163Dy (clone REA279) | Miltenyi | Cat#130-122-323 |
| anti-human CCR9-164Dy (clone L053E8) | BioLegend | Cat#358902 |
| anti-human CCR6-165Ho (clone G034E3) | BioLegend | Cat#353402 |
| anti-human NKG2D-167Er (clone REA797) | Miltenyi | Cat#120-014-229 |
| anti-human ITGB7-169Tm (clone REA441) | Miltenyi | Cat#130-122-309 |
| anti-human CD11b-172Yb (clone M1/70) | BioLegend | Cat#101214 |
| anti-human CXCR3-173Yb (clone REA232) | Miltenyi | Cat#130-108-022 |
| anti-human CD29-175Lu (clone TS2/16) | BioLegend | Cat#303021 |
| **Chemicals, peptides, and recombinant proteins** |  |  |
| ViaStain™ AOPI Staining Solution | Nexcelom | Cat#CS2-0106 |
| RPMI 1640 + glutamine | Corning | Cat#10-104-CV |
| pen strep | Gibco | Cat#15140122 |
| HEPES (1M) | Gibco | Cat#15630-080 |
| 0.5M EDTA | invitrogen | Cat#AM9260G |
| Hank's Balanced Salt Solution (HBSS) without Ca2+ Mg2+ | Thermo Scientific | Cat#88284 |
| HBSS with Ca2+ Mg2+ | Gibco | Cat#14025-076 |
| DNAse I | Sigma-Aldrich | Cat#10104159001 |
| Collagenase IV | Sigma-Aldrich | Cat#C4-BIOC |
| Maxpar® Cell Staining Buffer | Fluidigm | Cat#201068 |
| Maxpar® 10x Barcode Perm Buffer | Fluidigm | Cat#201057 |
| Heparin | Sigma |  |
| Formaldehyde, 37% | Electron Microscopy Sciences | Cat#15686 |
| PBS | Standard Vendors | N/A |
| Fetal Bovine Serum | Gibco | Cat#26140079 |
| RBC lysis buffer | G-Biosciences | cat# 786-672 |
| CryoStor CS10 freeze media | StemCell Technologies | Cat# 07930 |
| RNAlater | Sigma-Aldrich | Cat# R0901 |
| **Critical commercial assays** |  |  |
| eBioscience™ Fixable Viability Dye eFluor™ 506 | Invitrogen | Cat#50246097 |
| eBioscience™ 7-AAD Viability Staining Solution | Invitrogen | Cat# 00-6993-50 |
| EQ™ Four Element Calibration Beads | Fluidigm | Cat# 201078 |
| Dead cell removal kit | Miltenyi | Cat#130-090-101 |
| EasySep Human CD34 Selection Kit II | StemCell Technologies | Cat# 17856 |
| Chromium Next GEM Single Cell 3' Kit v3.1 | 10x Genomics | Cat#1000269 |
| Chromium Next GEM Chip G Single Cell Kit | 10x Genomics | Cat#1000127 |
| Dual Index Kit TT Set A | 10x Genomics | Cat#1000215 |
| Library Construction Kit | 10x Genomics | Cat#1000190 |
| Bioanalyzer High Sensitivity DNA Analysis Kit | Agilent | Cat#5067-4626 |
| **Deposited data** |  |  |
| Raw and analyzed single cell RNA-seq data |  |  |
| **Experimental models: Organisms/strains** |  |  |
| Mouse: NOD.Cg-Prkdcscid Il2rgtm1Wjl/SzJ (NSG), 8-9 weeks, Female | The Jackson Laboratory | Strain #005557 |
| **Software and algorithms** |  |  |
| Cytobank | Beckman Coulter | <https://www.beckman.com/flow-cytometry/software/cytobank-premium> |
| Helios software | Fluidigm |  |
| Matlab | MathWorks | <https://www.mathworks.com/products/matlab.html> |
| Matlab debarcoding tool |  |  |
| Prism 10 | GraphPad Software | <https://www.graphpad.com/> |
| Cell Ranger v6.1.1 | 10x Genomics |  |
| R version 4.2.2 (2022-10-31) | R Core |  |
| Seurat v4.3.0 | Hao and Hao et al. (2021) | <https://satijalab.org/seurat/>, 10.1016/j.cell.2021.04.048 |
| Python v3.11.3 |  | <https://www.python.org/downloads/release/python-3113/> |
| coreSC@ |  | <https://github.com/ChoBioLab/coreSC> |
| ImmunoSEQ Analyzer versions 2.0 and 3.0 | Adaptive Biotechnologies | <https://clients.adaptivebiotech.com/> |
| **Other** |  |  |
| CPT Tubes | BD | Cat#362753 |
| Cellometer Auto 2000 Cell Viability Counter | Nexcelcom | N/A |
| Cellometer disposable cell counting chamber slides | Nexcelcom | Cat#CHT4-PD100-002 |
| 70µm strainer | Fisherbrand | Cat#22-363-548 |
| 0.1µm spin filter | MilliporeSigma | Cat#UFC30VV00 |
| Attune NxT Flow Cytometer | ThermoFisher | N/A |
| EasySep Magnet | StemCell Technologies | Cat#18000 |
| MACS LS Columns | Miltenyi | Cat#130-042-401 |
| MACS MultiStand Separator | Miltenyi | Cat#130-042-303 |
| CyTOF2 Mass Cytometer | Standard Biotools | N/A |
| SuperSample fluidics system | Victorian Airships | N/A |
| Chromium iX instrument | 10x Genomics | cat#1000328 |
| Novaseq S4 | Illumina | N/A |
| NovaSeq 6000 Sequencing System | Illumina | N/A |
