## Supplementary figures and images for "The reparative immunologic consequences of stem cell transplantation as a cellular therapy for refractory Crohn’s disease"

### Supplementary Figure 1

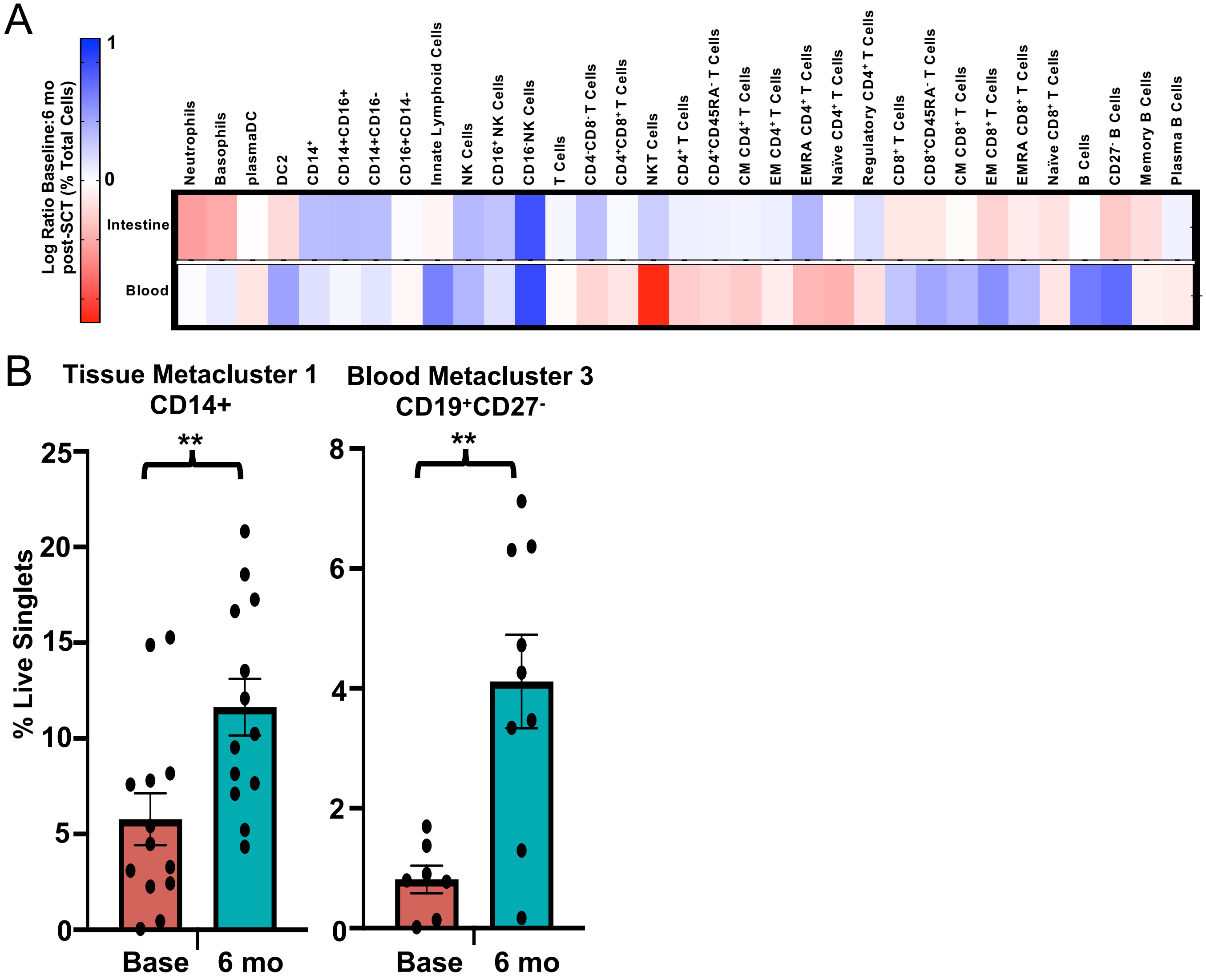

### Supplementary Figure 2

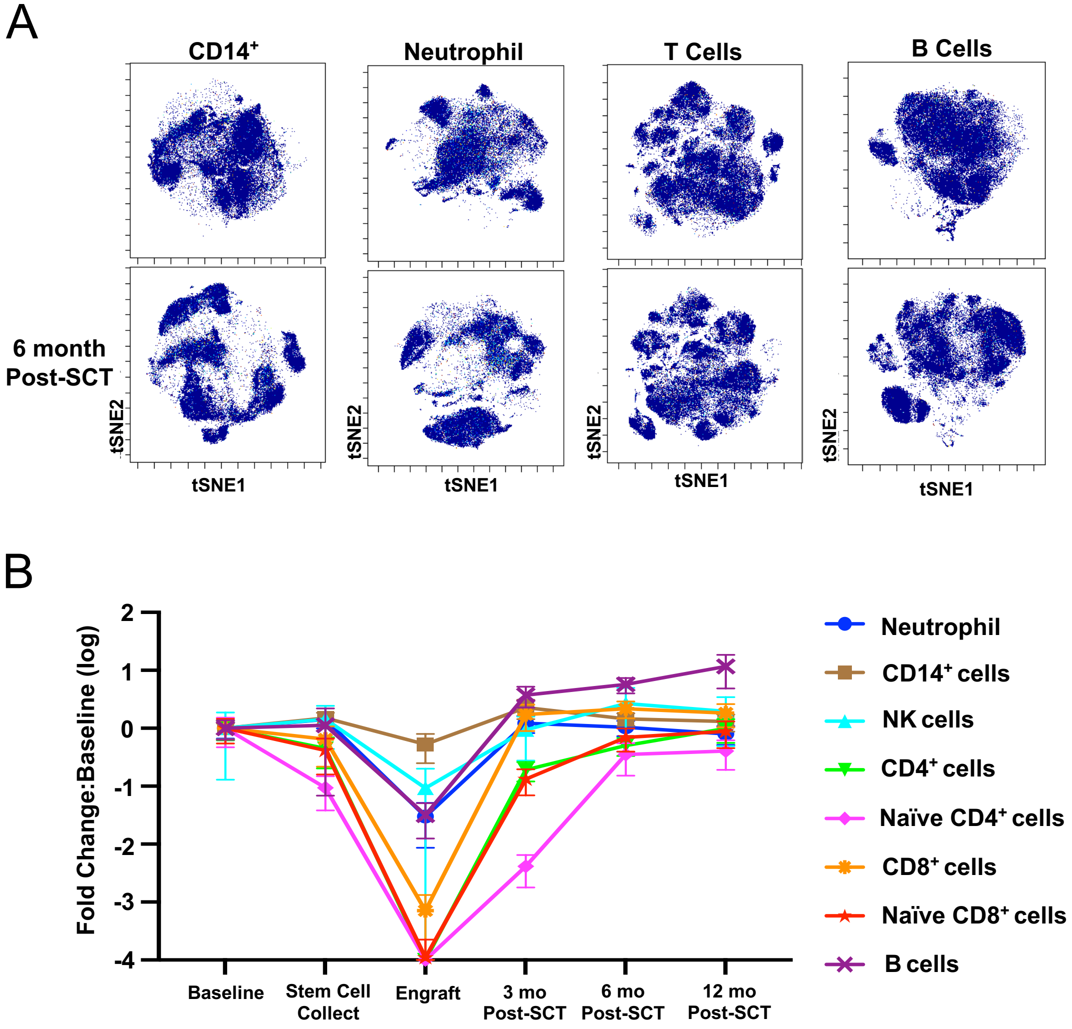

### Supplementary Figure 3

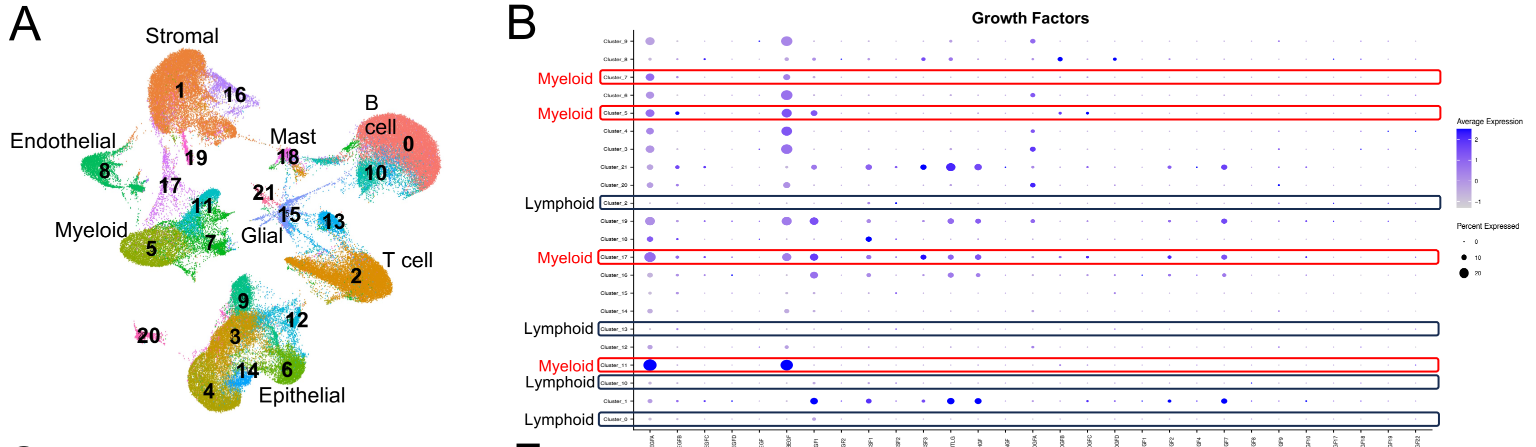


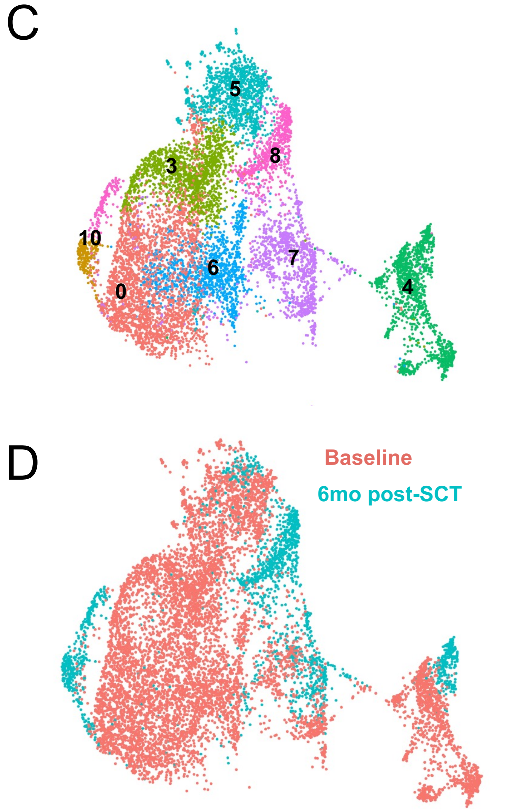


E

| Cluster | Baseline % | 6 mo post-SCT % |
|---|---|---|
| 0 | 32 | 3 |
| 3 | 18 | 1 |
| 4 | 14 | 13 |
| 5 | 14 | 9 |
| 6 | 14 | 2 |
| 7 | 8 | 23 |
| 8 | 1 | 37 |
| 10 | 0.1 | 11 |

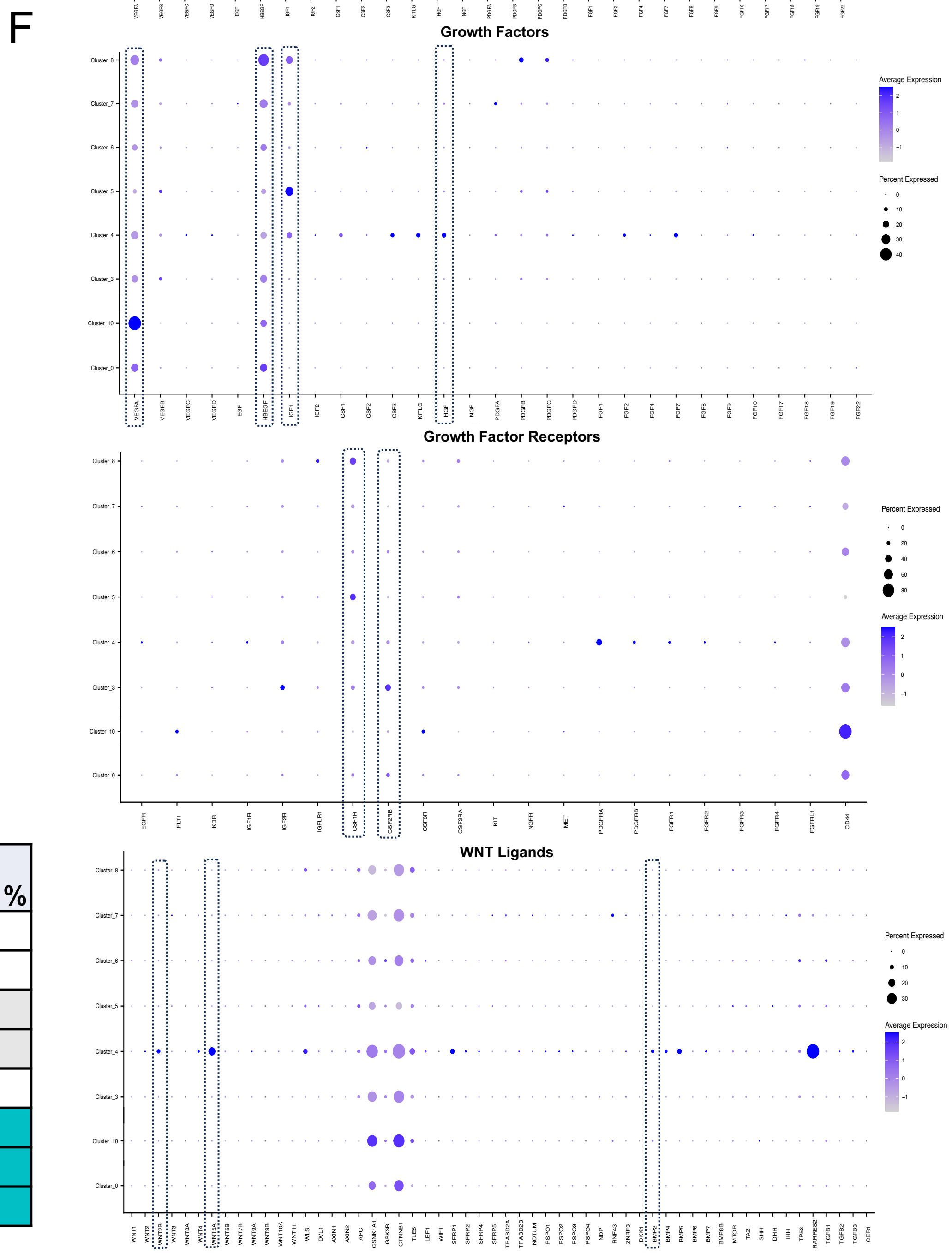

### Supplementary Figure 5

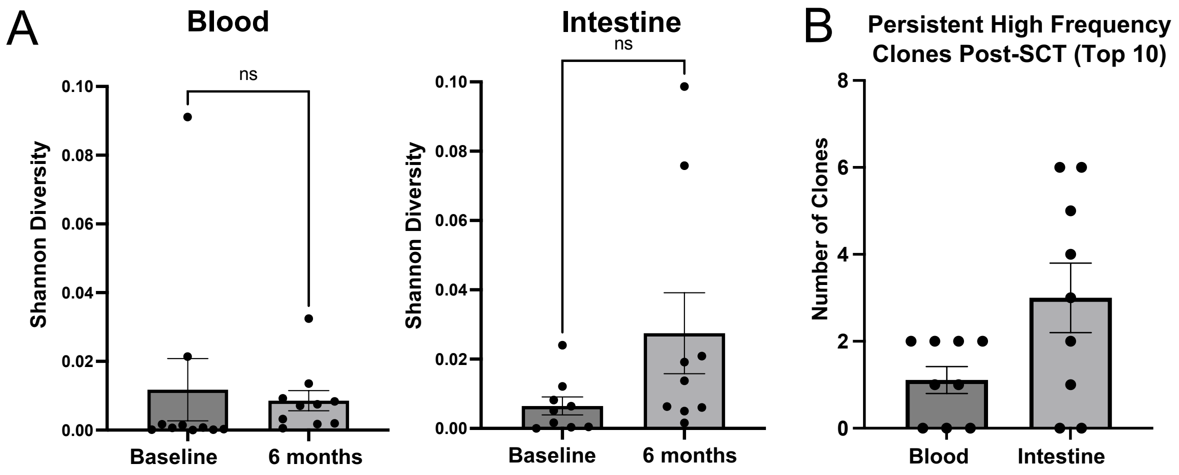

### Supplementary Figure 6

A

| | Pt 1 | Pt 2 | Pt 3 | Pt 4 | Pt 5 | Pt 6 | Pt 7 | Pt 8 | Pt 9 |
|---|---|---|---|---|---|---|---|---|---|
| **LT-HSC** | 4.77% | 0.26% | 0.00% | 7.11% | 0.04% | 1.61% | 2.48% | 0.40% | 5.56% |
| **IT-HSC** | 1.71% | 2.82% | 2.54% | 4.44% | 0.05% | 2.31% | 0.32% | 0.33% | 0.23% |
| **MPP/ST-HSC** | **54.67%** | **73.08%** | **53.99%** | **74.58%** | **85.25%** | **76.88%** | 3.01% | 3.55% | 1.62% |
| **Lineage Commit** | 25.34% | 12.82% | 40.94% | 1.23% | 1.21% | 2.51% | **89.86%** | **94.46%** | **83.80%** |

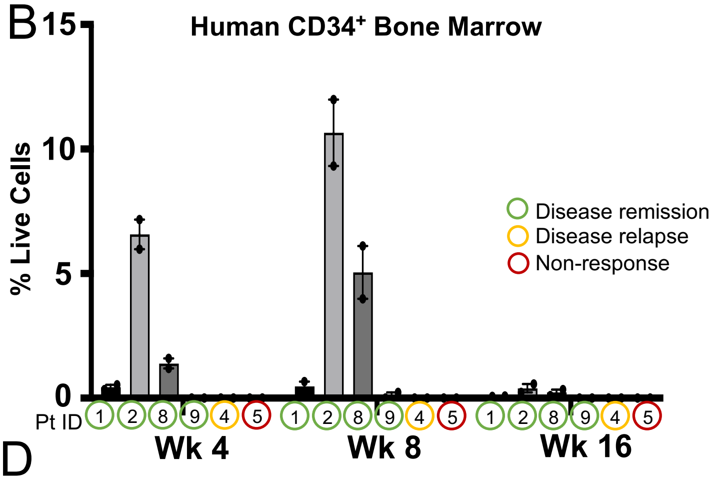


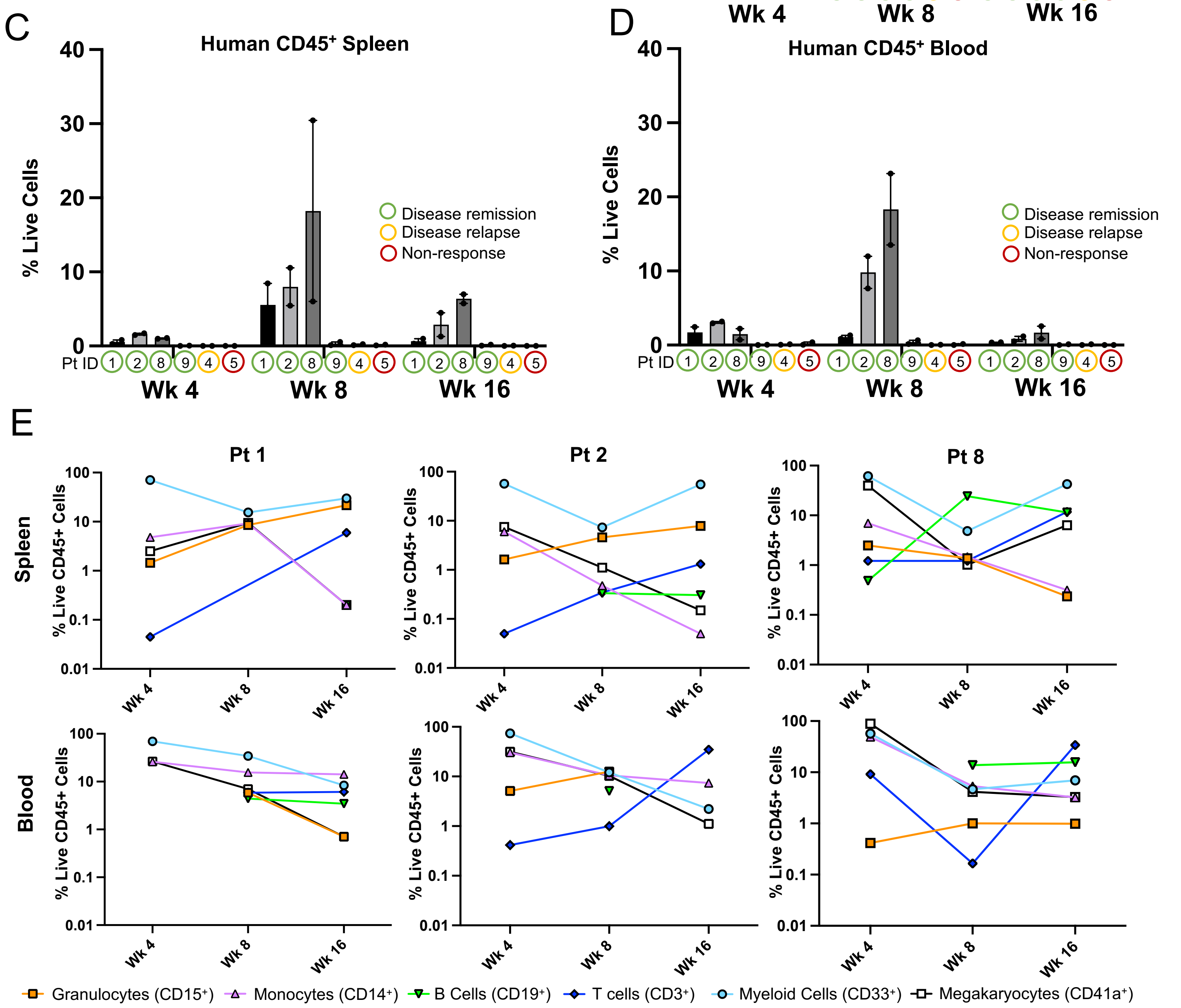

### Supplementary Figure 7

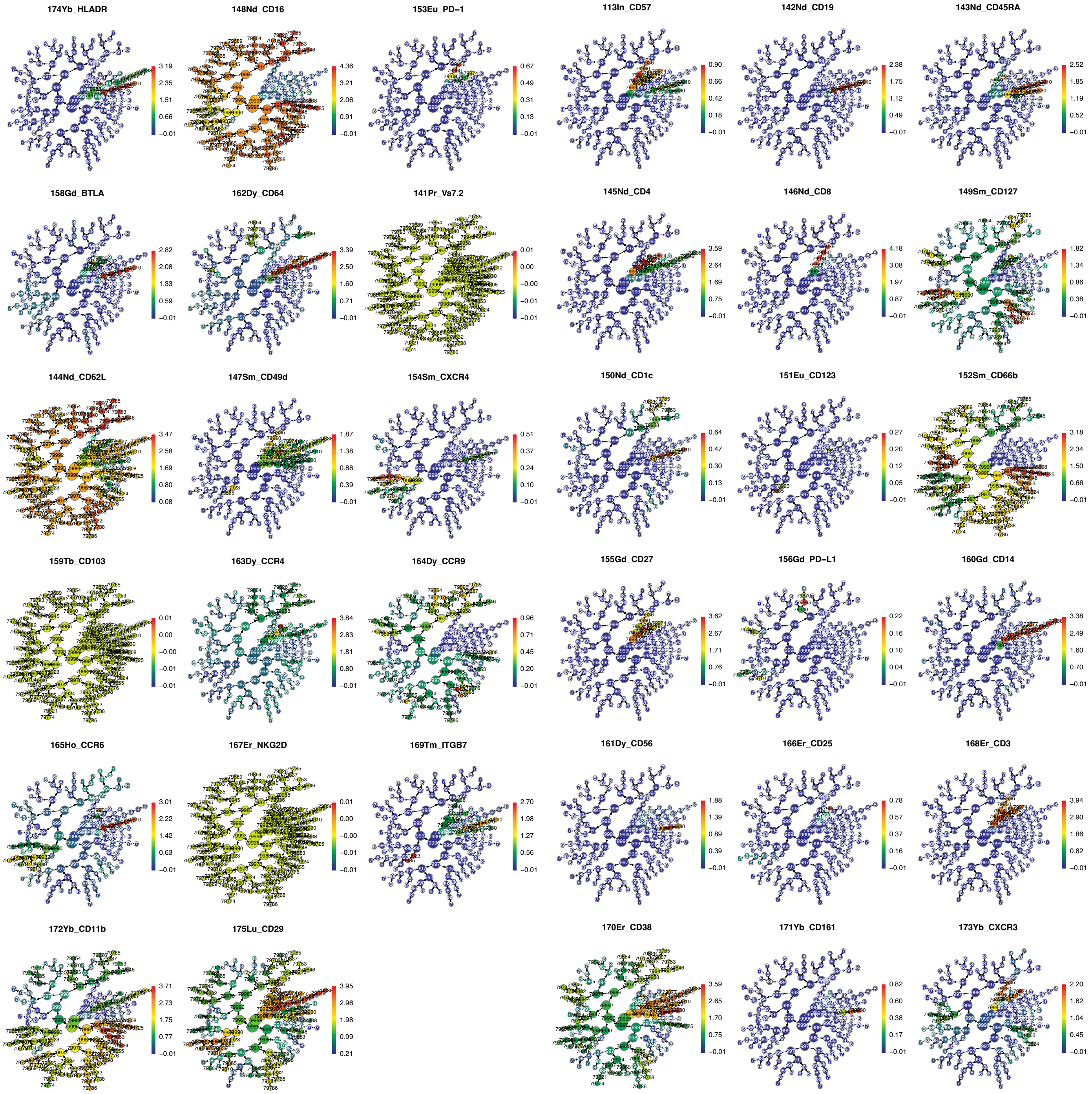

### Supplementary Figure 8

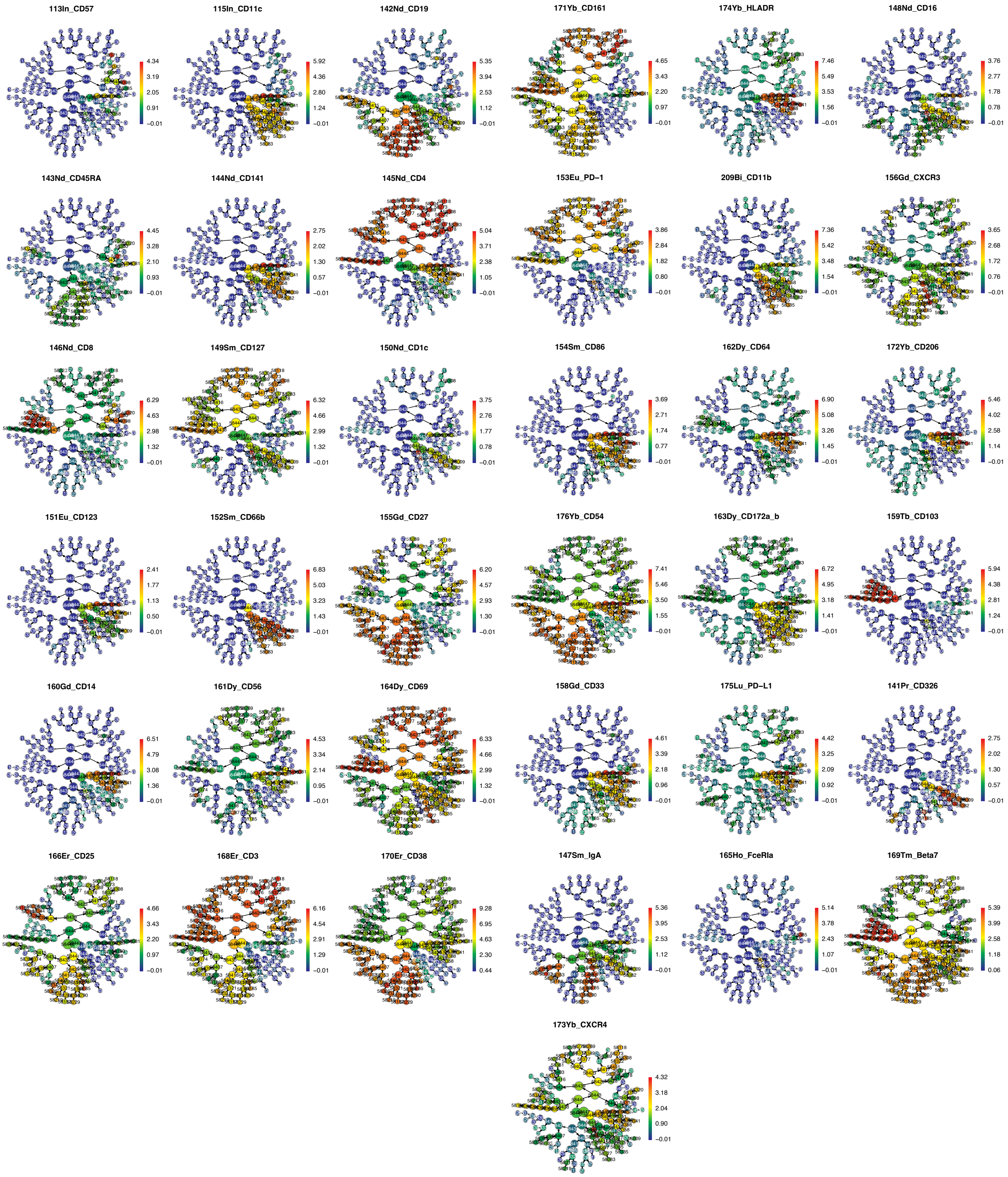

### Supplementary Figure 9

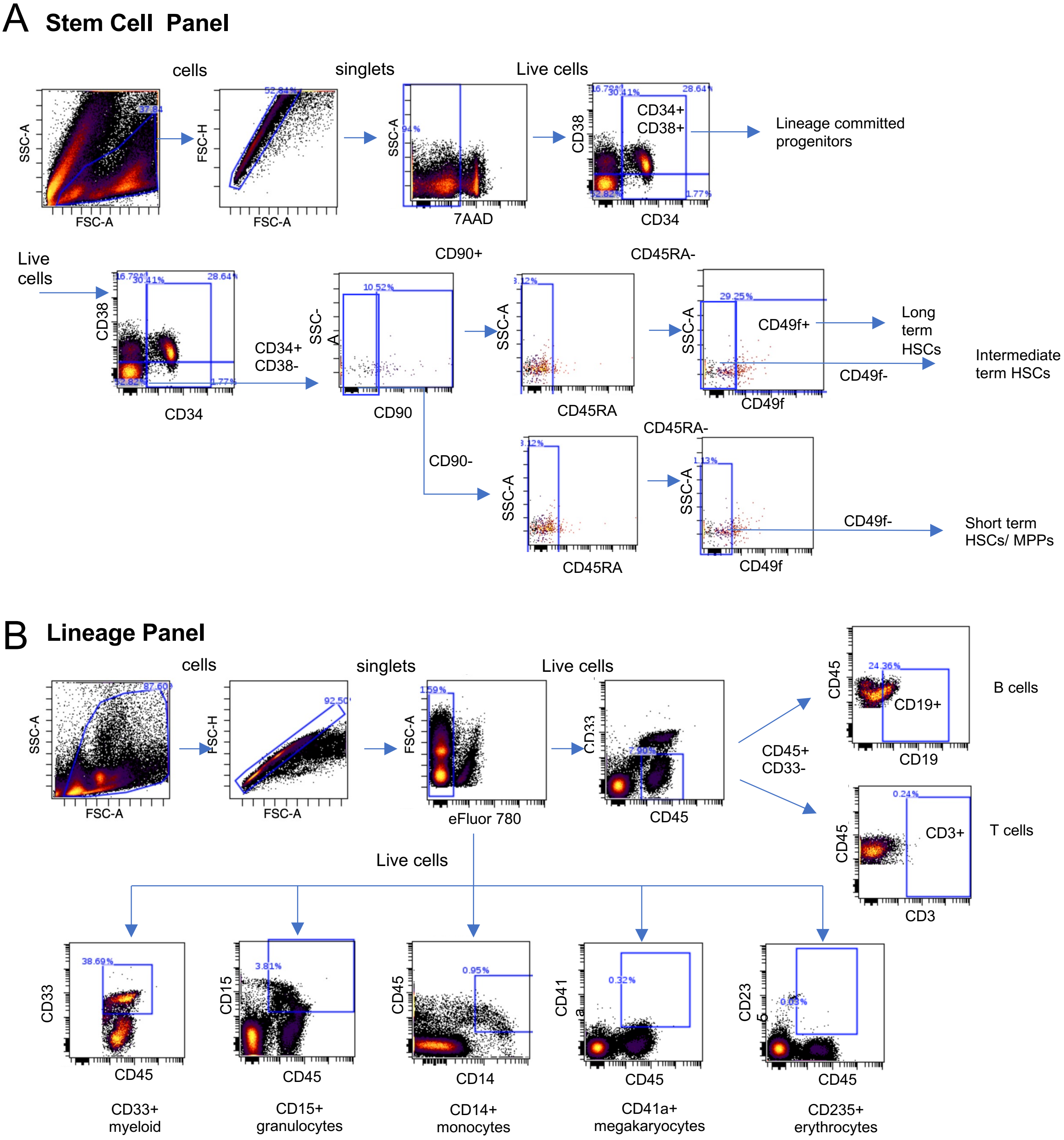
