## Supplementary Figure 4 for "The reparative immunologic consequences of stem cell transplantation as a cellular therapy for refractory Crohn’s disease"

A

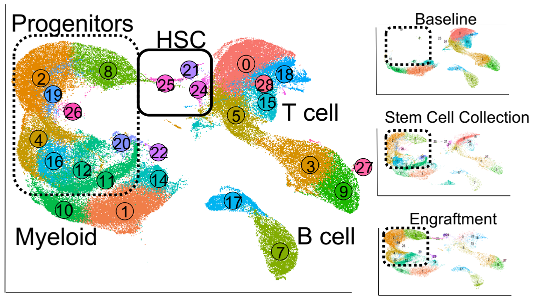


B

| Cluster | Annotation | Baseline | SCC | Engraft | 3mo | 6mo |
|---|---|---|---|---|---|---|
| 21 | HSC | 0 | 0.03 | 2.49 | 0 | 0.01 |
| 24 | HSC | 0.07 | 1.72 | 0.24 | 0.04 | 0.04 |
| 25 | HSC | 0.01 | 0.12 | 1.19 | 0 | 0 |
| 22 | Progenitor | 0 | 0 | 2.37 | 0 | 0 |
| 20 | Progenitor | 0.01 | 3.45 | 0.37 | 0 | 0 |
| 8 | Progenitor | 0.05 | 4.57 | 13.04 | 0.08 | 0.04 |
| 2 | Progenitor | 0 | 13.16 | 25.65 | 0 | 0 |
| 19 | Progenitor | 0 | 2.56 | 1.40 | 0 | 0 |
| 4 | Progenitor | 0.15 | 9.32 | 12.63 | 0.21 | 0.05 |
| 26 | Progenitor | 0 | 0.85 | 0.02 | 0 | 0 |
| 16 | Progenitor | 0.02 | 13.07 | 0.46 | 0 | 0 |
| 11 | Progenitor | 0.61 | 17.33 | 0.75 | 0.61 | 0.50 |
| 12 | Progenitor | 0 | 1.55 | 13.65 | 0 | 0 |
| 10 | Myeloid | 6.66 | 2.91 | 2.54 | 11.09 | 2.53 |
| 1 | Myeloid | 15.48 | 6.66 | 6.83 | 25.04 | 7.60 |
| 14 | Myeloid | 5.96 | 1.88 | 1.69 | 4.87 | 4.47 |
| 27 | NK cells | 0 | 0.16 | 0.46 | 0 | 0 |
| 3 | NK cells | 9.49 | 2.41 | 3.31 | 9.42 | 13.58 |
| 9 | NK cells | 6.31 | 1.89 | 2.79 | 6.04 | 10.06 |
| 7 | B cells | 4.83 | 0.93 | 2.67 | 10.26 | 19.08 |
| 17 | B Cell | 2.75 | 0.70 | 1.68 | 5.00 | 10.73 |
| 0 | T cell | 27.89 | 10.42 | 0.63 | 4.10 | 4.73 |
| 18 | T cell | 2.06 | 0.09 | 1.31 | 10.75 | 12.04 |
| 28 | T cell | 0 | 0.69 | 0.01 | 0 | 0 |
| 5 | T cell | 8.88 | 2.55 | 1.42 | 11.11 | 12.08 |
| 15 | T cell | 8.78 | 0.97 | 0.40 | 1.37 | 2.45 |

0 % Total cells 25

C

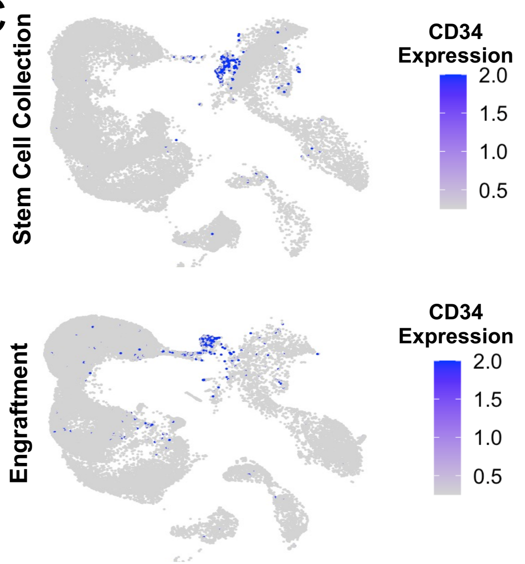


D

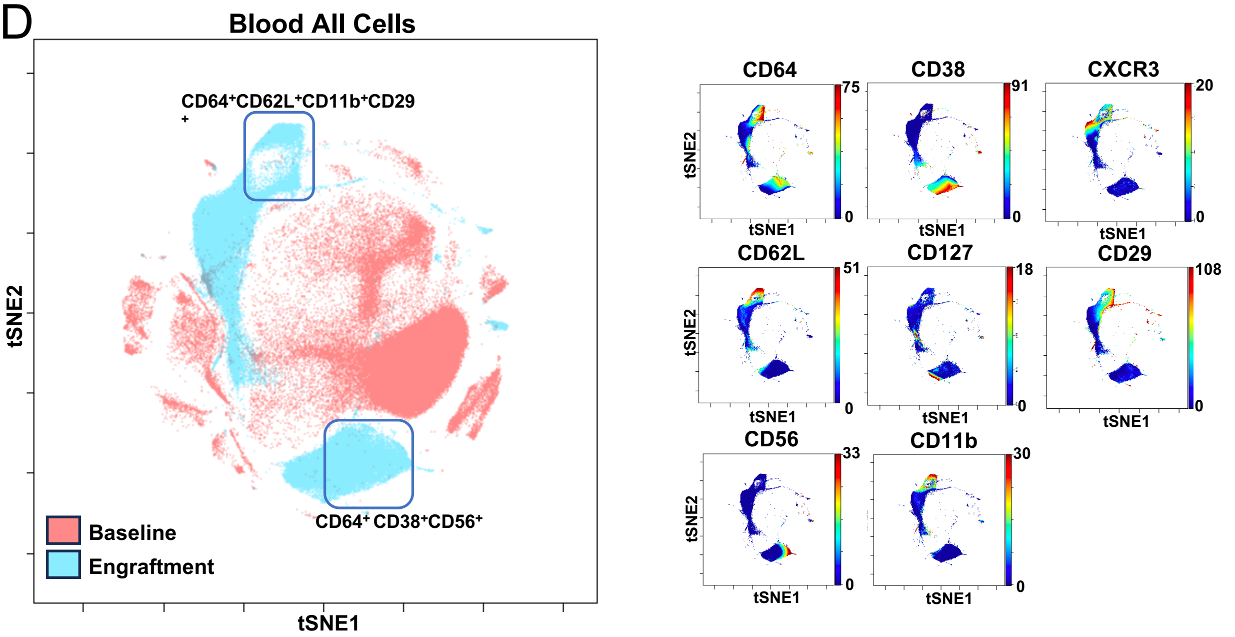
